# Selection of a Promiscuous Minimalist cAMP Phosphodiesterase from a Library of *De Novo* Designed Proteins

**DOI:** 10.1101/2023.02.13.528392

**Authors:** J. David Schnettler, Michael S. Wang, Maximilian Gantz, Christina Karas, Florian Hollfelder, Michael H. Hecht

## Abstract

The ability of unevolved amino acid sequences to become biological catalysts was key to the emergence of life on Earth. However, billions of years of evolution separate complex modern enzymes from their simpler early ancestors. To study how unevolved sequences can develop new functions, we screened for enzymatic activity in a collection of > 1 million novel sequences based on a *de novo* 4-helix bundle library of semi-random sequences. To mirror evolutionary selection for biological function, we screened the collection using ultrahigh-throughput droplet microfluidics to identify features that yield phosphoesterase activity. Characterization of active hits demonstrated that acquiring new function required a large jump in sequence space: screening enriched for truncations that removed > 40% of the protein chain and introduced a catalytically important cysteine. The truncated protein dimerized into a dynamic α-helical structure, consistent with the idea that gain of function was accompanied by an increase in structural dynamics relative to the parental 4-helix bundle. The purified protein catalyzes the hydrolysis of a range of phosphodiesters, with the greatest activity toward the biological second messenger cyclic AMP (cAMP). The novel cAMPase is a manganese-dependent metalloenzyme and catalyzes cAMP hydrolysis with a rate acceleration on the order of 10^9^ and catalytic proficiency on the order of 10^14^ M^−1^, comparable to large enzymes shaped by billions of years of evolution. These findings suggest that fragmentation to modular primordial peptides can be a fertile avenue for introducing structural and functional diversity into proteins.

**GRAPHICAL ABSTRACT:** 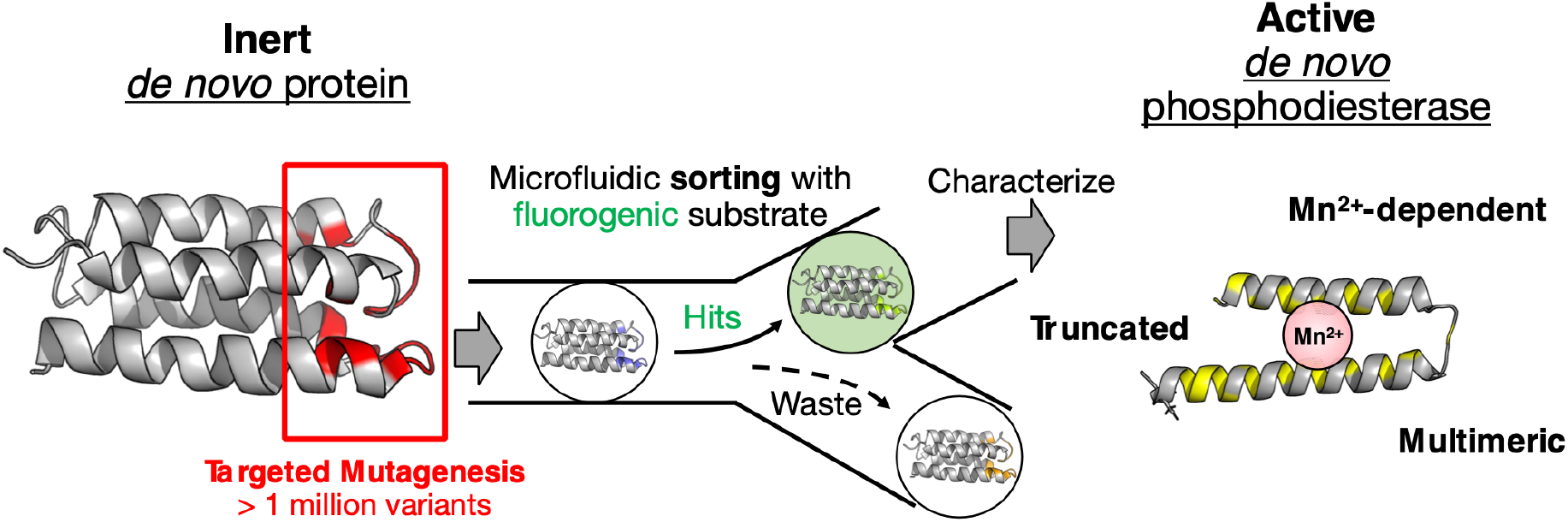

## INTRODUCTION

Living systems depend upon chemical reactions that, if uncatalyzed, occur too slowly to support life. Therefore, evolution has selected biological catalysts – enzymes – that speed up reactions to rates sufficient to sustain survival and growth. The enzymes found in present day proteomes are typically well-ordered finely tuned proteins, which provide rate enhancements far superior to catalysts created by design. Moreover, most enzymes in the current biosphere are relatively large, with bacterial and eukaryotic proteins having average lengths of 320 and 472 amino acids, respectively^1^.

It is relatively straightforward to envision how these large and highly active enzymes evolved by iterative selection for improvements of progenitor sequences that were already active and folded. However, it is far more challenging to understand how the first minimally active sequences arose *de novo* from inactive random polypeptides. Understanding how functional enzymes emerged from random sequences is particularly challenging in light of a large body of work showing that fully random sequences (i) rarely fold into well-ordered soluble structures^2,3^, and (ii) rarely bind biologically relevant small molecules. For example, a seminal study by Keefe and Szostak^4^ showed that random libraries of 80-residue polypeptides include sequences that bind ATP at a frequency of approximately one in 10^11^. Catalysis is an even greater challenge than binding. The promiscuity of catalysts has been invoked to facilitate emergence of new reactions after gene duplication and adaptive evolution of a side activity^5,6^. Metagenomic libraries have been found to contain catalysts for ‘unseen’, non-natural reactions with a xenobiotic substrate^7^ at a frequency of one in 10^5^. But eliciting function from non-catalytic sequences instead of refunctionalizing an active enzyme is a much more difficult proposition, with very few examples on record^8,9^. Computational design has been successful in principle^10^, but is far from routine and creating high efficiency catalysts rivalling the efficiency of evolved enzymes has so far been elusive. At an extreme, creationists – arguing that divine intervention is necessary – have estimated that sequences capable of forming modern day enzymes might be extremely rare^11^: as low as one in 10^50^–10^74^.

These considerations generate a tension between the abundant success of life on earth and the extreme rarity of sequences capable of sustaining such life in the ‘vastness of sequence space’, because the step between inactive and active sequences (postulated in the Dayhoff hypothesis^12,13^ as the historical link between small peptides and contemporary proteins) seems a near-unsurmountable hurdle. To probe this contradiction, we set out to explore whether biologically relevant catalysts can be isolated from collections of unevolved *de novo* sequences. Because we anticipated active catalysts would be rare and fully random sequences rarely fold into stable soluble structures^2,3^, several features were incorporated into the screen to enhance the likelihood of success. (i) As a starting scaffold we chose a collapsed globular structure, known to fold into a stable 4-helix bundle^14^. This scaffold S-824 is a 102-residue protein (previously isolated from a semi-random library^15^) without natural function or known activity; (ii) Ultrahigh-throughput screening in microfluidic droplets at a rate of ≈ 0.8 kHz was used to search libraries generated from S-824 for rare hits amidst a background of inactive sequences in a 1.7 million-membered library; (iii) The screened substrates contained both phosphodiesters and phosphotriesters, thereby broadening the type of catalysts that might be discovered. (iv) Finally, a mixture of typical divalent metals (often found as cofactors in hydrolases) was added to the screen, thereby allowing the isolation of *de novo* metalloenzymes.

Our screen yielded a truncated, structurally dynamic 59-residue enzyme able to bring about rate accelerations from ≈ 10^9^ in efficient promiscuous turnover of phosphodiesters in the presence of manganese, including the unreactive substrates second messenger cyclic AMP (cAMP). The rapid identification of a simple yet functional short protein with considerable activity provides support for the Dayhoff hypothesis^12,13^ and illustrates a scenario rapid conversion of a non-catalytic peptide into a proficient catalyst that is crucial for evolution.

## RESULTS

### Microfluidic screening of a library of *de novo* sequences yields catalytically active proteins

S-824, a stable 102-residue-long 4-helix bundle protein, was randomized in its apical loops and helix termini to create an active site cavity surrounded by catalytic groups and producing a library of ≈ 1.7 million variants (**Figure 1a**)^16^. The starting scaffold of S-824 in turn had previously been isolated from a library randomized with binary patterning of polar and nonpolar amino acids^15,17^, its 3-dimensional structure is known (PDB ID: 1p68)^14^ and there are no natural homologs and no evolved function known. Such a 4-helix bundle scaffold tolerates enormous sequence diversity, with a hydrophobic core as only minimal constraint^17^, minimizing the chances of sacrificing the folded structure and thus allowing us to interrogate the functional potential of diverse randomized sequences.

**Figure 1:**
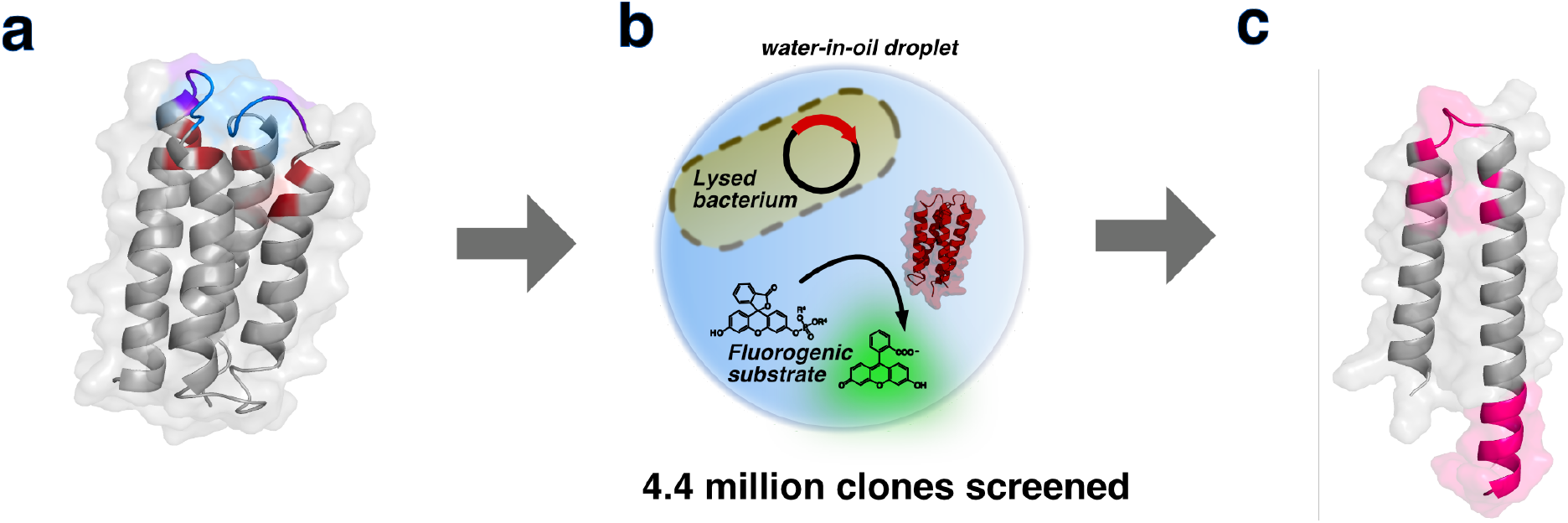
Droplet screening of a library of *de novo* designed 4-helix bundles enriches truncated sequences with phosphoesterase activity. **(a)** *De novo 4-helix bundle library*. A library^16^ containing ≈ 1.7 million variants based on the stably folded *de novo* designed 4-helix bundle protein S-824 was screened for phosphoesterase activity. The diversified residues of S-824 are shown in red (degenerate codon: NDT) and blue (degenerate codon: VRC). **(b)** *Ultrahigh-throughput microdroplet screening*. The library was subjected to fluorescence-activated droplet sorting (FADS) on a microfluidic chip, and the 0.1–0.2% most fluorescent droplets (of a total of 4.4 million screened) were selected. **(c)** *Truncated peptide with phosphoesterase activity*. The selection yielded catalytically active, truncated peptides consisting of a helix-turn-helix of ≈ 60 amino acids (mutated sites shown in magenta), illustrated here with a structural model of mini-cAMPase generated with AlphaFold2/ColabFold^21,22^.

The parental S-824 sequence showed no phosphoesterase activity in our screening assay, so to maximize the chances of capturing active enzymes we incorporated diversity into all components of the screen: not only were the sequences diverse, but we also screened for different catalytic activities simultaneously by using a bait mixture of fluorogenic phospho-di- and -triester substrates (**Ext. Data Fig. 1**). Moreover, because many natural phosphoesterases utilize divalent metal cofactors, we increased the chance of finding a hit by supplementing the reaction with a mixture of MnCl_2_, ZnCl_2_, and CaCl_2_. We sampled all these variables combinatorially in an ultrahigh-throughput manner by using microfluidic droplets, which allow screening of the entire library in a single experiment. In this microfluidic assay, individual bacterial cells expressing a single library sequence were co-compartmentalized with the mixture of fluorogenic substrates and metals, and then lysed in the droplet (**Figure 1b, Figure S1**)^18,19^. After incubation, the droplets were dielectrophoretically sorted at ultrahigh throughput by fluorescence-activated droplet sorting (FADS) on a microfluidic chip^20^. Sorted hits were subsequently identified by recovering and sequencing their plasmid DNA.

For the initial sort, we chose permissive conditions to maximize the number of screened clones and enhance the likelihood of finding hits. In total, 10.3 million droplets (corresponding to ≈4.4 million clones) were screened in 4 hours, and the 0.2 to 0.5% most fluorescent droplets were selected. The collected hits were clonally expanded and re-sorted under more stringent conditions to further enrich catalytically active clones. After this cumulative enrichment by droplet screening, ≈ 250 clones were arbitrarily picked for secondary screening in microtiter plates. Although the starting protein S-824 had very low phosphoesterase activity above background, average activity levels increased with each round of sorting (**Figure S2)**, indicating that screening cumulatively enriched sequences with phosphate hydrolase activity.

### Enrichment of truncations along with gain of function

Sequence analysis of these single hits from the secondary screen revealed that some of the most active sequences (12/14) had frameshift mutations which produced truncated protein sequences (**Figure 1c**). Indeed, Next-Generation Sequencing (NGS) analysis of the entire library before and after screening (≈ 1.4 million unique variants) suggested that droplet screening had enriched single nucleotide deletions along with the gain of function. These deletions led to frameshift mutations and consequently an enrichment in premature stop codons (from 17% to 27%; **Figure 2a** and **Ext. Data Fig. 2**). This observed enrichment stands in stark contrast to the fact that intra-gene stop codons are typically strongly selected against, as they usually lead to a loss of structure and function^23–25^. Thus, the paradox of an observed enrichment of truncated variants – as a general trend across the library – indicated that truncations might contribute to the selected phosphoesterase activity. Further analysis of their positional distribution revealed a high abundance of truncations specifically at positions 40 and 60, with 18% of all sequences being truncated at one of these positions after the second sorting round. Truncations at both positions are enriched compared to the input library (1.2 and 2.8-fold, **Figure 2b**). These truncated sequences lack the two C-terminal helices of the 4-helix bundle (**Figure 1c**) and form the template for a shorter helix-loop-helix motif.

**Figure 2:**
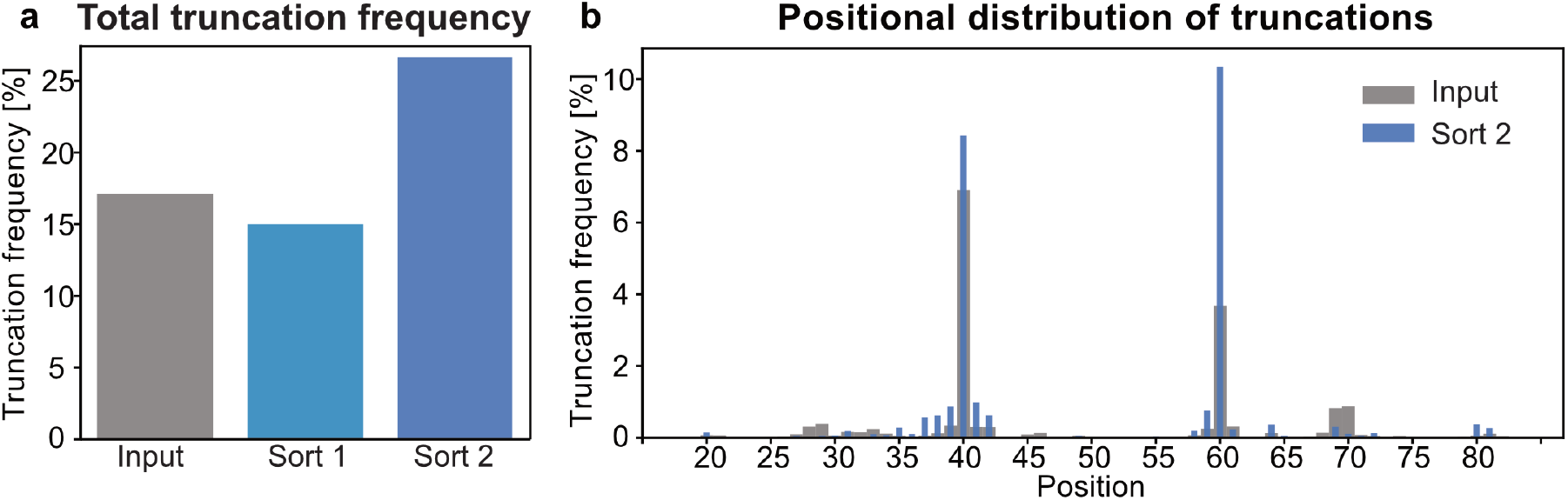
Next-Generation Sequence analysis reveals enrichment of truncated proteins across the library. **(a)** Relative frequency (percentage of truncated sequences of total number of sequences) of truncated reads in input library (grey; 17%), after sorting 1 (light blue; 15%) and sorting 2 (dark blue; 27%) reveals 1.6-fold enrichment of premature stop codons after sort 2 albeit close to 0% would be expected **(b)** Frequency of truncations at every sequenced position in input library (grey, broad bars) and after sorting 2 (blue, narrow bars) shows 1.2 and 2.8-fold enrichments of truncations at position 40 and 60, respectively, after sorting 2.

### A manganese-dependent *de novo* phosphodiesterase

We focused on the analysis of one clone (subsequently named mini-cAMPase, see below) with high catalytic activity, strong expression and good solubility and disentangled the combinatorial elements of the screen (multiple substrates; multiple metal cofactors). Manganese was required for activity (**Figure 3a**) while not altering thermal stability (**Ext. Data Fig. 3**), consistent with a catalytic rather than structural role. However, manganese binding could not be measured by Isothermal Titration Calorimetry (ITC, **Figure S4**), which prevented determination of affinity and stoichiometry.

**Figure 3:**
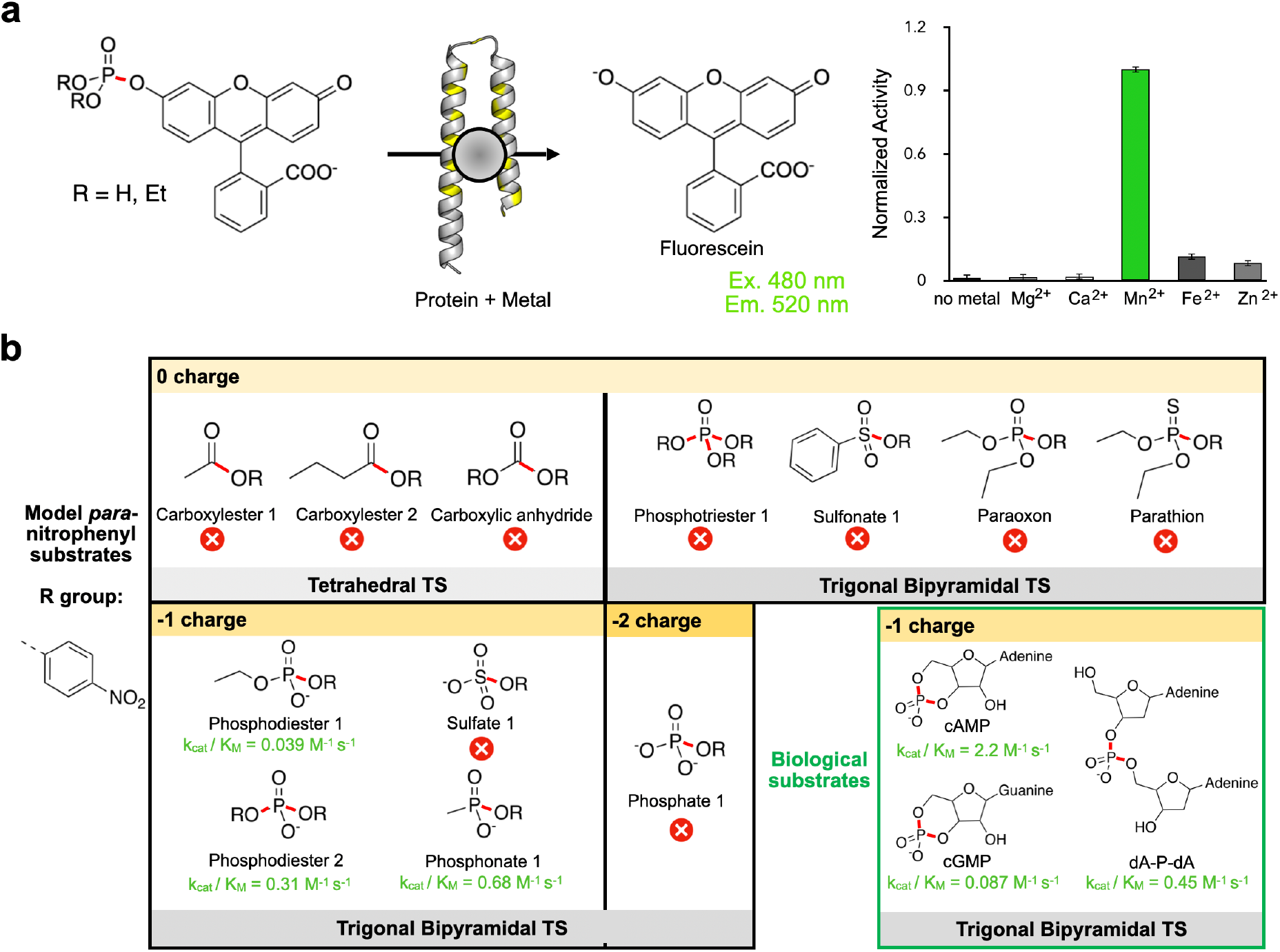
Metal requirement and substrate scope. **(a)** Metal dependence of the enzymatic reaction for the mixture of fluorogenic bait substrates. 100 μM of mini-cAMPase was incubated with 200 μM substrate mixture and 200 μM of either Ca^2+^, Mg^2+^, Mn^2+^, Fe^2+^ or Zn^2+^. The histogram shows mini-cAMPase requires Mn^2+^ for catalytic activity. **(b)** Activity was tested against substrates sampling different ground state charges (from 0 to −2) and transition state (TS) geometries (tetrahedral and trigonal-bipyramidal). mini-cAMPase hydrolyses phosphodiesters **2** and a phosphonate **1**, with the highest activity being towards cAMP. The scissile bond is highlighted in red. Red cross symbols indicate no detectable activity.

Next, we probed the substrate specificity by challenging the novel enzyme with a range of model substrates containing *p*-nitrophenyl leaving groups and encompassing a range of ground state charges and transition state geometries. As shown in **Figure 3b**, these experiments revealed the novel enzyme catalyzes hydrolysis of phosphodiester and phosphonate substrates carrying a single negative charge. Furthermore, activity was limited to substrates hydrolyzed through a trigonal-bipyramidal transition state.

To assess activity beyond model compounds with nitrophenyl leaving groups, we extended the scope of substrates to biologically important phosphodiesters. In contrast to the expectation that promiscuous activities identified in screens for thermodynamically undemanding substrates cannot easily be extended to more difficult reactions, we found the novel enzyme catalyzes the hydrolysis of less reactive biological phosphodiester substrates, cAMP and cGMP. It also shows some activity toward deoxyadenosine dinucleotide (dA-P-dA), which can be considered a simple model for DNAse activity. Because our novel protein is most active towards cAMP, we named it mini-cAMPase.

The observation of catalytic activity for a *de novo* protein expressed in *E. coli* inevitably raises concerns about the possibility of contaminating activity from endogenous proteins^26,27^. We ruled out this possibility by performing several control experiments, showing that the observed activity cannot be attributed to endogenous *E. coli* cAMPase (CpdA)^28^ or other contaminating proteins: **(i)** mini-cAMPase requires a different metal cofactor and has a much lower Michaelis constant than endogenous *E. coli* cAMPase. **(ii)** The enzymatic activity co-purifies with the mini-cAMPase fraction, independent of the purification method. **(iii)** Sequence changes in mini-cAMPase correlate with changes in enzymatic activity. These control experiment thus provide evidence that mini-cAMPase is a *de novo* phosphodiesterase (for details, see **Note 1.2 in the Supporting Information** and **Figures 4a–c** and **Ext. Data Fig. 4 and 5**).

**Figure 4:**
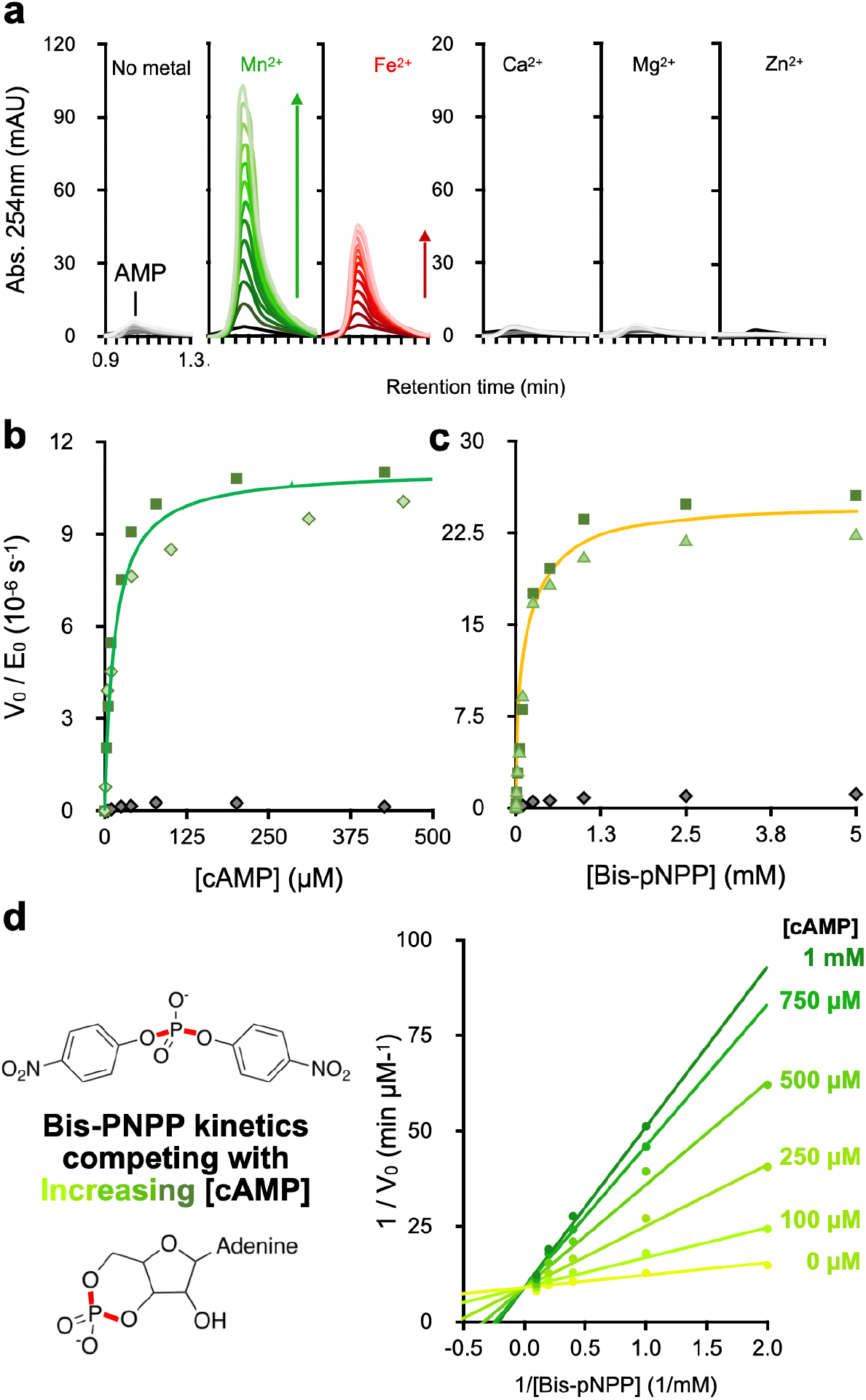
Kinetic characterization of enzymatic activity. (**a**) Metal-dependence of mini-cAMPase activity. Shown is the raw appearance of AMP on RP-HPLC when 100 μM of mini-cAMPase is incubated with 100 μM of the respective divalent metal and 250 μM cAMP; each successive trace is 1.5 h apart, with a total time of 18 h. (**b**) Michaelis-Menten kinetics for cAMPase activity in biological duplicate with 50 μM protein and 200 μM MnCl_2_, measured at 25 °C in phosphate buffered saline (pH 7.4). The purification control (grey) shows the background activity for the same preparation steps on the protein without the His_6_-tag. (**c)** Michaelis-Menten plot for phosphodiesterase activity of 50 μM mini-cAMPase and 200 μM MnCl_2_ with the model substrate bis(*p*-nitrophenyl) phosphate in biological duplicate. Each color shows a biological replicate. As before, the purification control is shown in grey. (**d**) Competition between bis-pNPP and cAMP hydrolysis shown by Lineweaver-Burke plot. Lines connect kinetics for a single concentration of cAMP, with higher cAMP concentrations shown in darkening hues of green. The shared y-intercept indicates competitive inhibition with *K_i_* = 70 ± 8 μM cAMP. Michaelis-Menten kinetics were made for bis(*p*-nitrophenyl) phosphate hydrolysis by 100 μM protein with 200 μM MnCl_2_ in the presence of increasing concentrations of cAMP.

**Figure 5:**
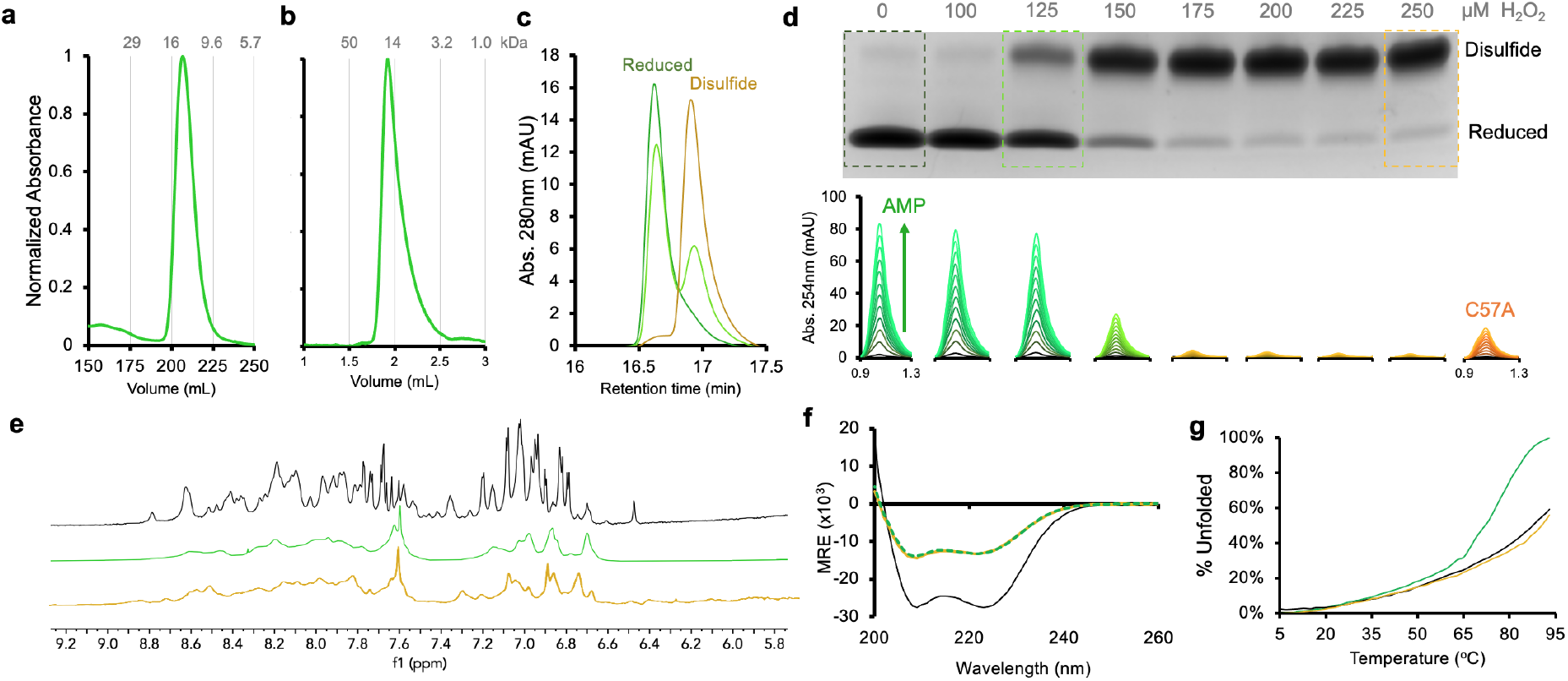
Mini-cAMPase is a dynamic α-helical dimer. Across all panels, yellow represents the disulfide-bonded dimer, green is the reduced protein, and black is the parental protein, S-824. **(a)** Preparation-scale and (**b**) analytical-scale size-exclusion chromatography show reduced mini-cAMPase elutes at double its monomeric molecular mass (i.e., it is a noncovalent dimer). **(c)** Reverse-phase HPLC traces show reduced (green), partly oxidized (yellow-green, 125 μM H_2_O_2_), and fully oxidized (yellow, 250 μM H_2_O_2_) protein. **(d)** Activity under a gradient of increasingly oxidizing conditions to promote disulfide formation. The gel is an SDS-PAGE without reducing agent, showing the relative abundance of disulfides. Below each gel lane is the corresponding activity, of 50 μM mini-cAMPase incubated with 200 μM Mn^2+^ and 250 μM cAMP, shown as the appearance of AMP (assayed by RP-HPLC) measured every 2 h for 24 h. The mutant C57A is included for comparison. **(e)** ^1^H NMR showing the amide region for S-824 (black) and mini-cAMPase (reduced green, oxidized yellow). Broad and poorly resolved peaks indicate the protein does not form a well-ordered structure and remains dynamic even after oxidation. **(f)** Circular dichroism (CD) spectra show that mini-cAMPase (green, yellow) is helical, but less so than the parent 4-helix bundle (black). There is no discernible shift in secondary structure caused by oxidation of C57. MRE: mean residue ellipticity. (**g**) Melting curves measuring CD at 222 nm show that oxidized mini-cAMPase (yellow, T_M_ > 90 °C) has a melting curve similar to S-824 (black, T_M_ > 90 °C) with both proteins more thermostable than the reduced mini-cAMPase (green, T_M_ ≈ 72 °C).

### Kinetic characterization and demonstration of a common active site

Kinetic characterization of mini-cAMPase with both model and biological substrates found the largest catalytic efficiency (*k_ca_*_t_/*K_M_*) with cAMP, revealing a catalytic efficiency of 2.2 M^−1^s^−1^ which corresponds to a rate acceleration of ≈ 7 × 10^9^ and a catalytic proficiency of ≈ 7 × 10^14^ M^−1^ (**Table 1**, **Figure 4b,c**). Because of this level of activity, and because of its biological relevance, cAMP became the focus of further characterization. As with the fluorogenic substrate mixture, we found that mini-cAMPase best hydrolyzes cAMP in the presence of manganese, although it has some residual activity in the presence of iron (**Figure 4a**). The Michaelis-Menten kinetics show slow turnover, but a definite *V_max_*, suggesting active site saturation for both cAMP and bis-*p*NPP. Notably, the parental sequence S-824 lacks detectable cAMPase activity (**Ext. Data Fig. 6f** and **8b**) but shows low activity towards bis-*p*NPP (*k_cat_*/*K_M_* ≈ 4 × 10^−3^ M^−1^ s^−1^, so ≈ 80-fold lower than mini-cAMPase, **Ext. Data Table 1** and **Ext. Data Fig. 8c**).

**Table 1:** **Michaelis-Menten constants with phosphonate and phosphodiester substrates** measured at 50 μM enzyme concentration in 50 mM HEPES-NaOH, 150 mM NaCl, 200 μM MnCl_2_, pH 8.0 at 25 °C. Since mini-cAMPase functions as a dimer, kinetic parameters are calculated per protein dimer. The indicated error is the error of the non-linear fit.

| Substrate | $k_{cat}$<br>($10^{-6} \text{ s}^{-1}$) | $K_M$<br>($\mu\text{M}$) | $k_{cat}/K_M$<br>($\text{M}^{-1}\text{s}^{-1}$) | $k_{cat}/k_{uncat}$ | $(k_{cat}/K_M)/k_{uncat}$<br>($\text{M}^{-1}$) |
| --- | --- | --- | --- | --- | --- |
| *p*-nitrophenyl-methylphosphonate <sup>a</sup> | $17 \pm 0.2$ | $25 \pm 3$ | 0.68 | $1 \times 10^6$ | $4 \times 10^{10}$ |
| *p*-nitrophenyl-ethyl-phosphate <sup>b</sup> | $8.0 \pm 0.2$ | $210 \pm 31$ | 0.039 | $1 \times 10^{10}$ | $6 \times 10^{13}$ |
| bis(*p*-nitrophenyl-)phosphate <sup>b</sup> | $50 \pm 4$ | $160 \pm 40$ | 0.31 | $7 \times 10^{10}$ | $5 \times 10^{14}$ |
| cAMP <sup>c</sup> | $22 \pm 4$ | $10 \pm 3$ | 2.2 | $7 \times 10^9$ | $7 \times 10^{14}$ |
| cGMP <sup>c</sup> | $19 \pm 0.4$ | $210 \pm 10$ | 0.087 | $6 \times 10^9$ | $3 \times 10^{13}$ |
| dA-P-dA <sup>b</sup> | $0.91 \pm 0.1$ | $20 \pm 4$ | 0.045 | $1 \times 10^9$ | $6 \times 10^{13}$ |

<sup>a</sup> Based on $k_{uncat} = 1.7 \times 10^{-11} \text{ s}^{-1}$ for *p*-nitrophenyl phenylphosphonate at pH 7.5 and 30 °C<sup>29</sup>.
<sup>b</sup> Based on $k_{uncat} = 7 \times 10^{-16} \text{ s}^{-1}$ for P-O cleavage in dineopentyl phosphate at pH 7 and 25 °C<sup>30</sup>.
<sup>c</sup> Based on $k_{uncat} = 3 \times 10^{-15} \text{ s}^{-1}$ for P-O cleavage in ethylene phosphate (a cyclic phosphodiester) at pH 7 and 25 °C<sup>31</sup>. For further details, see **Note 1.6 in the Supplementary Information**.

While both the model substrates and the biological substrates (**Figure 3b**) are phosphodiesters, these compounds have very different structures. Therefore, we were curious about whether mini-cAMPase hydrolyzed these substrates using the same active site. To address this question, we probed whether hydrolysis of the model chromogenic substrate bis-*p*NPP could be inhibited by cAMP. As shown in **Figure 4d** (and elaborated in **Figure S5**), hydrolysis of bis-*p*NPP in the presence of increasing concentrations of cAMP showed competitive inhibition with a *K_i_* of 70 ± 8 μM for cAMP. These results indicate that despite their structural and electronic differences, both substrates rely upon the same active site.

### Mutational and structural analysis of mini-cAMPase

Mini-cAMPase carries both truncations and substitutions as compared to S-824. To address the relative contributions of both factors to the observed catalytic activity, we tested two alternative constructs: to probe the importance of the truncation, we incorporated the substitutions observed in mini-cAMPase – but *not* the truncation – into the full-length parental sequence S-824, creating a sequence dubbed ‘Substituted-824’. On the other hand, to probe the role of the substitutions, we introduced the truncation – but not the substitutions – into the parental protein, producing ‘Short-824’ (**Ext. Data Fig. 7a**). Testing the expression and activity of these constructs indicated that the designed substitutions and the unexpected truncation might potentially *both* contribute to cAMPase activity (**Ext. Data Fig. 7b)**. Short-824 expresses very poorly (**Ext. Data Fig. 7c**), suggesting that one role of the library substitutions is to stabilize the truncated protein. Substituted-824 expressed well and, in contrast to the parental sequence S-824, shows activity towards both phosphodiester substrates, albeit with a reduced *k_cat_* (≈ 7-fold towards cAMP and ≈ 17-fold towards bis-*p*NPP as compared to mini-cAMPase, **Ext. Data Fig. 7d)**. This indicates that both the substitutions and the truncations contribute to catalysis: the enzymatic activity is largely bestowed by the substitutions introduced by mini-cAMPase, and potentially to a minor part by the truncation. However, it should be noted that Substituted-824 carries five substitutions towards the C-terminal end which neither mini-cAMPase nor S-824 carry and whose effect on catalytic activity is unknown.

To identify the residues directly involved in catalysis or binding of the Mn^2+^ co-factor, we mutated several residues individually to alanine and measured Michaelis-Menten kinetics of the mutants (**Ext. Data Table 1**). We targeted residues that might be involved in manganese binding, including histidine and cysteine, as well as the arginine consistently introduced by the truncations (**Ext. Data Fig. 8a**). All introduced mutations reduced *k_cat_*, with C57A having the largest impact, lowering *k_cat_* ≈ 6.5-fold towards cAMP and ≈ 3.7-fold towards bis-*p*NPP (**Ext. Data Fig. 8b,c**). This indicates that C57A is involved in but not essential to catalysis.

Because only two of the original four helices are present, we surmised that mini-cAMPase might form a helical hairpin that dimerizes into a 4-helix bundle. Consistent with this expectation, size-exclusion chromatography (SEC) shows that mini-cAMPase elutes as a dimer (**Figure 5a,b**). This dimer was formed under reducing conditions and confirmed by LC/MS to be fully reduced at its only cysteine residue C57 (**Ext. Data Fig. 9**). Further experiments under oxidizing conditions showed that disulfide bridge formation also leads to dimerization but abolishes activity completely (**Figure 5c,d and Ext. Data Fig. 9**), implicating that the disufide-linked dimer structure is incompatible with efficient catalysis. In an alternative scenario, the observed loss of activity could be due to concomitant further oxidation of the Mn^2+^ co-factor under the oxidizing conditions.

^1^H-NMR, and circular dichroism (CD) spectroscopy further indicate that mini-cAMPase is structurally less defined than its parental sequence S-824 (**Figure 5e–g**), at least in the absence of manganese. This could indicate that it dynamically samples an ensemble of states, potentially including conformations that enable activity. Alternatively, in an induced fit model, metal binding could potentially induce a more defined, functional structure in mini-cAMPase which is not sampled by the catalytically inactive ancestor S-824.

## DISCUSSION

Mini-cAMPase has several notable features: **(i)** It was isolated from a library of variants that was relatively small (≈ 1.7 × 10^6^) compared to the vast sequence space available to natural evolution (20^102^ ≈ 10^132^ for a protein of the length of S-824). **(ii)** Although the parental S-824 shows no phosphoesterase activity, the selected mini-cAMPase recruits a metal co-factor to catalyze phosphodiester hydrolysis with high catalytic proficiency. **(iii)** While the inactive parental sequence was 102 residues long, screening for activity led to a truncation protein of approximately half the length of the parent. **(iv)** Although the S-824 parent is a well-ordered monomeric 4-helix bundle, the selected mini-cAMPase enzyme forms a 2×2 helix dimer with additional dynamic potential after loss of a covalent constraint.

The catalytic efficiency (*k*_*ca*t_/*K_M_*) of mini-cAMPase (2.2 M^−1^s^−1^; **Table 1**) for cAMP is only ≈ 1000-fold lower than, e.g., cAMPase (*CpdA*) from *E. coli* (**Table S3**). Furthermore, although the catalytic efficiency of mini-cAMPase is well below that of the ‘average’ natural enzyme acting on its *preferred* substrate (*k*_*ca*t_/*K_M_* ≈ 10^5^ M^−1^s^−1^)^32,33^, it is only ≈ 14-fold lower that the median value of 31 M^−1^s^−1^ reported for natural enzymes catalyzing *promiscuous* reactions^34^. This implicates that, like naturally occurring promiscuous enzymes, mini-cAMPase could be optimized to high efficiency in few rounds of directed evolution. Importantly, mini-cAMPase displays a substantial rate enhancement (*k_cat_/k_uncat_*) on the order of 10^9^ and a catalytic proficiency (*(k_cat_/K_M_)/k_uncat_*) in the range of ≈ 10^14^ M^−1^ (Table 1). This can be compared with natural enzymes, where rate enhancements and catalytic proficiency vary widely^35^ with rate enhancements ranging from 10^5^ to 10^17^ and catalytic proficiencies ranging from ≈ 10^8^ to 10^23^. Further, it surpasses the rate enhancement of a zinc-based biomimetic phosphodiesterase model by 5 orders of magnitude^36^. Thus, although turnover is slow because the substrate is thermodynamically challenging, the actual catalytic effect of mini-cAMPase approaches that of some large natural enzymes. However, the turnover rates of naturally evolved phosphodiesterases specifically acting on cAMP range from 10^−1^ to 10^3^ s^−1^ (**Table S3**), with rate enhancements in the range of 10^14^ to 10^17^, thus still surpassing mini-cAMPase by several orders of magnitude.

mini-cAMPase is a rare example of rapid functionalization of a short, inactive peptide, validating the Dayhoff hypothesis and dismissing the skeptical (or creationist) view that sequence space does not hold sufficient solutions for catalytic challenges from peptides with no biological ancestry. Here we show how catalysis of a difficult biologically relevant reaction with accelerations (*k_cat_/k_uncat_*) approaching those of large natural enzymes selected by billions of years of evolution can be brought about in an efficient screen of million-membered library that nevertheless represents only a small fraction of all possible diversity.

Intriguingly, the recruitment of a divalent metal cofactor for phosphodiester hydrolysis by mini-cAMPase recapitulates the evolution of naturally occurring phosphodiesterases. Extant, naturally occurring cAMP-hydrolysing enzymes have emerged independently in three different folds, converging towards mechanisms that involve a divalent metal ion cofactor^37^. Most natural phosphodiesterases use Zn^2+^, and the *E. coli* cAMPase (CpdA) requires Fe^2+^ or Mg^2+^. In contrast, mini-cAMPase requires Mn^2+^ for activity, suggesting an unprecedented variation to the theme of divalent metal ion catalysis.

The strategy of starting a combinatorial screen by randomizing a stable fold followed by ultrahigh-throughput screening (offering multiple cofactors and substrates) led to mini-cAMPase, which has a catalytic proficiency surpassing that of designed (and subsequently randomized and selected) peptides by several orders of magnitude. For example, a partly designed and further evolved 4-helix bundle metalloprotein with a catalytic proficiency of 9.3 × 10^10^ M^−1^ for carboxyesters^38^, and further designed and evolved into a Diels-Alderase^39^ with a catalytic proficiency of 2.9 × 10^11^ M^−1^. Similarly, a previously reported^40^ *de novo* helix-turn-helix peptide that dimerizes into a 4-helix bundle is able to hydrolyze the phosphodiester *p*-nitrophenyl phosphate (in absence of metal) with a second-order rate constant of 1.58 × 10^−4^ M^−1^ s^−1^, resulting in a catalytic proficiency of ≈ 2 × 10^11^ M^−1^, which is three orders of magnitude lower than mini-cAMPase.

The isolation of mini-cAMPase from a relatively small library (compared to the theoretical size of sequence space) was facilitated by two factors: (i) First, the use of microfluidic droplet sorting allowed screening for multiple-turnover catalysis at very high throughput (4.4 million clones screened in just 4 hours), ensuring that most clones of our million-membered library were sampled at least once (≈ 2.6-fold oversampling). Likewise, previous examples of generating function from inactive scaffolds also relied on ultrahigh-throughput screening directly for catalysis, like mRNA display^8,41^. (ii) Second, targeted randomization of a *de novo* sequence designed by binary patterning to fold into a stable 4-helix bundle structure, enhanced the number of sequences in the library displaying a stable fold.

Unexpectedly, selection for functional catalysts enriched library members which ‘escape’ the pre-defined stable 4-helix bundle fold by truncation to specific lengths, forming a helix-turn-helix motif. While such major truncations are usually expected to be detrimental to protein function, Next-Generation Sequencing confirmed this as a general trend across the entire library (**Figure 2**). As characterized in detail for the case of mini-cAMPase, the remaining 59 residues maintain the binary patterning of the original design. The observation of dimer formed of α-helices (**Figure 5a,b**), suggests a return to a 4-helix bundle fold, albeit with additional degrees of freedom that may tap dynamic effects in binding and catalysis (**Figure 5e–g**).

The selection of a truncated library member invites speculation that the 4-helix bundle fold of the library ancestor S-824 is very stable but structurally limited. Compared to its parent S-824, the structure of mini-cAMPase is highly dynamic (**Figure 5e–g**), possibly fluctuating between multiple conformational states and/or dimeric arrangements. This feature might represent a departure from S-824’s ‘frozen’ conformation, sampling arrangements conducive to catalytic function by conformational selection^42–44^. Although structural disorder has been linked to ineffective catalysis^45,46^, the sampling of new conformational states may also provide functional innovation that can be further enriched in subsequent steps of evolution^47,48^ or simply be signs of damage incurred by mutation^47,48^. The disorder seen in mini-cAMPase can thus be likened to a catalytically proficient molten globule enzyme^49^ or the melting of a zinc finger that conferred ligase activity to a non-catalytic scaffold^9^.

Evolved natural enzymes typically fold into large well-ordered structures with pre-organized active sites. Presumably, these were selected to favor highly active and substrate-specific ‘specialists’ that can be allosterically regulated. In the early stages of evolution, however, it may have been advantageous to express dynamic sequences that sampled an ensemble of states. While these molten structures likely had low activities, they may have been able to catalyze several chemical reactions, acting on a range of substrates. Such multifunctional ‘generalists’ would have enhanced the “catalytic versatility of an ancestral cell that functioned with limited enzyme resources”^5,50^. In addition, small, dynamic proteins may have advantages as starting points for further evolution and adaptation. While large modern enzymes are restricted by an epistatic burden that causes mutations to interfere with structural and functional innovation^51–54^, small dynamic proteins could escape the limits imposed by cooperative effects and become functionally more versatile and more evolvable by reducing the cost of innovation.

Going beyond making Dayhoff’s hypothesis more plausible, the case of mini-cAMPase suggests that proficiency can be generated by going back to the origins of life, in smaller, less structurally defined folds and that targeted randomization of closer-to-primordial scaffolds, when analyzed by ultrahigh-throughput technologies, provides unexpected catalytic solutions (e.g., truncation) that complement the lessons about the acquisition of function in contemporary enzymes.

## METHODS

### Library preparation

The library encoding the partly randomised *de novo* designed gene S-824 was cloned from the original pET11a vector^16^ into the high-copy number vector pASK-IBA5+ (IBA Life Sciences, Germany) using the restriction sites XbaI and BamHI. This allowed highly efficient DNA recovery by transformation after droplet sorting as well as strain-independent tetracycline-inducible expression^55^.

### Microfluidic library screening

Microfluidic library screening was carried out as previously described^56^. In brief, the library was electroporated into *Escherichia coli* cells (E. cloni 10G Elite; Lucigen, USA), yielding ≈ 10^7^ colonies after overnight incubation on agar plates, as determined by serial dilution. After induction and incubation for protein expression, cells were washed and encapsulated together with the substrate and metal mixtures and lysis agent in monodisperse water-in-oil droplets on a flow-focusing chip (**Figure S1a**), generating droplets with a volume of 3 pL at rates of 0.5 to 3 kHz. The droplets were collected into a closed storage chamber, fabricated from an eppendorf tube as previously described^57^. The microfluidic devices were fabricated by soft lithography as previously described^56^. After incubation at room temperature), droplets were reinjected from the collection tubing into the sorting device (**Figure S1b**). In contrast to the previously described sorting device^56^, this chip featured an additional flow of “bias oil ” from the side which forced the droplets further away from the hit channel at a flow rate of 30 μL/h, reducing the number of false positive droplets accidentally flowing into the hit channel. Droplets were sorted according to their fluorescence at a rate of ≈ 0.8 kHz and collected into a tube pre-filled with water. After sorting, the hit droplets were de-emulsified by addition of 1H,1H,2H,2H-perfluorooctanol (Alfa Aesar, USA), the solution was purified and concentrated by column purification and the plasmid DNA was recovered by electroporation into highly electrocompetent *E. coli* cells as previously described in detail^56^. In the initial sorting, permissive conditions were chosen: cells were encapsulated at an average droplet occupancy (λ) of 0.43 and droplets were incubated for three days. Out of 10.3 million screened droplets, 53 000 droplets were sorted (≈ top 0.5%). In the second sort, screenings were conducted under more stringent conditions, with cell encapsulation at λ = 0.1 and droplets were incubated for seven days. Out of 4.4 million screened droplets, 7500 droplets were sorted (≈ top 0.2%).

### Microtiter plate screening

To quantify the lysate activity of library variants, individual colonies were picked and grown in 96-deep-well plates in 500 μL LB medium (containing 100 μL/mL carbenicillin) at 37 °C/1050 rpm for 14 h. Subsequently, 20 μL of overnight cultures were used to inoculate 880 μL of TB medium (containing 100 μL/mL carbenicillin and 2 mM MgCl_2_) for expression cultures in 96-well deep-well plates which were grown at 37 °C/1050 rpm for 2–3 h until OD_600_ ≈ 0.5. Expression was then induced with anhydrotetracycline (final concentration 200 ng/mL; IBA Life Sciences, Germany) and carried out for 16 h at 20 °C and 1050 rpm shaking. Cells were pelleted by centrifugation at 3200 rcf for 60 min and the supernatant was then discarded. Subsequently, the cells were lysed by a freeze-thaw cycle, followed by resuspension of the dry pellets by vortexing (≈1 min) and subsequent addition of 150 μL lysis buffer (HEPES buffer containing 0.35X BugBuster Protein Extraction Reagent (Novagen) and 0.1% Lysonase Bioprocessing Reagent (Novagen)). Cells were incubated for 20 min at room temperature in lysis buffer on a tabletop shaker (1000 rpm) and subsequently subjected to 30 min thermal denaturation at 75 °C. The lysate was cleared by centrifugation for 1 h at 3200 rcf and 60 μL of the supernatant were used for the activity assay. For the reaction, 140 μL of the bait substrate mixture were added to 60 μL aliquots of the cleared lysate in microtiter plates. The assay was carried out in HEPES buffer with a final concentration of 10 μM of the bivalent cation mixture (ZnCl_2_, MnCl_2_, CaCl_2_) and 20 μM of the fluorescein phosphate bait substrate mixture. The formation of fluorescein was recorded in a plate reader (Infinite M200; Tecan, Switzerland) for 30–60 min at an excitation wavelength of 480 nm and an emission wavelength of 520 nm.

### Next-Generation Sequencing and Data Analysis

After droplet sorting and DNA recovery by electroporation, plasmid DNA was extracted from all obtained *E. coli* colonies using the GeneJET Plasmid Miniprep Kit (ThermoFisher Scientific). The variable region of the library (position 19 to 83) was amplified with PCR using Q5 polymerase (NEBnext UltraII Q5 Master mix) with primers including adapters for NGS in the overhang and different indices for each sample for multiplexing (**Table S1**). The cycle number was optimized using qPCR with variable template concentrations to be below 15 cycles for the final PCR reaction to minimize amplification bias. AMPure Speed Beads (Beckman Coulter) were used to purify DNA after amplification. The samples were processed into Illumina TruSeq libraries by the University of Cambridge, Department of Biochemistry sequencing facility according to the manufacturer’s instructions. Sequencing was performed with one Illumina MiSeq 2 x 300 bp run (20% PhiX spike-in) yielding 8.5 × 10^6^ sequences for the input library, 4.8 × 10^5^ sequences for sorting 1 and 6.5 × 10^5^ sequences for sorting 2. Adapters were removed, reads were merged, and individual sequences were counted using the DiMSum pipeline^58^. The processed read counts were analyzed using custom python scripts to count the frequency of truncations and frameshift mutations among all library members and at individual positions.

### Protein Expression and Purification

A single colony of *E. coli* BL21 carrying the plasmid to express protein of interest was used to inoculate a starter culture in LB medium and grown at 37 °C for 12 h. This started culture was used to inoculate 1 L of LB medium (1:200 dilution), which was grown at 37 °C until it reached an optical density at 600 nm of 0.6, at which time anhydrotetracycline was added to a final concentration of 200 ng/mL to induce protein expression. After 12 h of expression at 18 °C, cells were harvested via centrifugation and stored at −80 °C.

To purify protein, the cells were resuspended in phosphate buffer (50 mM Na_2_HPO_4_, 300 mM NaCl, pH 8.0) and lysed by sonication on ice at 45% amplitude for twenty 10 s bursts with 50 s between sonication events. Following centrifugation and filtration, clarified lysate was loaded onto a nickel column (GE HisTrap HP, 5 mL volume) and purified by imidazole elution (phosphate buffer with 500 mM imidazole). His_6_-tagged protein was eluted with 375 mM imidazole, while untagged protein still weakly stuck to the column and could be eluted with 10 mM imidazole. This was followed by size exclusion chromatography (SEC, HiLoad 26/600 Superdex 75pg) with Tris buffer. The elution time of the peak by SEC corresponds to ≈ 14 kDa when compared to protein standards, which supports a dimeric structure. Mutations did not alter the SEC profile.

To keep protein reduced throughout, 1 mM DTT was added after elution from the nickel column, and 1mM TCEP was added after elution from the sizing column. Proteins were aliquoted, frozen in liquid nitrogen, and stored at −80 °C for subsequent assays. All protein was also purified metal-free (*apo*) by the addition of 5 mM EDTA (pH 8.0) after elution from the nickel column. EDTA was removed by the subsequent SEC step. For protein oxidation, reducing agent was removed by PD-10 desalting column (GE healthcare) exchange to Tris buffer at a final protein concentration of approx. 120 μM. The protein was then split into samples to which increasing concentrations of hydrogen peroxide were added to promote disulfide formation^59^. Samples were then left in the cold room (4 °C) in the dark for 24 h. In the absence of hydrogen peroxide, the protein remained reduced due to residual reducing agent, but increasing concentrations of hydrogen peroxide allowed the protein to form disulfide-bonded dimers (**Figure 7c–d**). To monitor the presence/formation of disulfide-bonded dimers, proteins were analyzed by HPLC as described below, or on an 12% SDS-PAGE (BioRad) at 20 μM concentration without added reducing agent in the loading buffer.

### Size exclusion Chromatography

Size exclusion chromatography (SEC) was performed on an ÄKTA Pure FPLC (General Electric). Preparation-scale SEC was done on a HiLoad 26/600 Superdex 75pg column, and analytical SEC was run on a 5/150GL Superdex 75 increase column (Cytiva). Calibration was done against external standards γ-globulin (bovine), Ovalbumin (chicken), Myoglobin (horse), and Vitamin B-12 (Bio-Rad).

### Confirmation of Protein Purity

To assay for contaminating endogenous *E. coli* proteins, both reverse-phase HPLC and mass spectrometry were employed (**Figure S6**). To validate protein purity by reverse-phase HPLC, 10μL of purified protein was injected to a C-18 column (Agilent Zorbax 300SB-C18, 5 μM, 2.1 x 150mm) without dilution or buffer exchange. Solvent A was water with 0.1% TFA, and solvent B was acetonitrile with 0.1% TFA. The gradient was 97% solvent A / 3% solvent B from 0 to 5 min, 97% solvent A / 3% B to 100% solvent B from 5 to 25 min to elute proteins, 100% solvent B from 25 to 30 min to wash the column, and then 97% solvent A / 3% solvent B from 30 to 35min to re-equilibrate. The only peak detected was the *de novo* protein. Moreover, fractions were then run on ESI-MS (Agilent 6210 TOF LC/MS) to confirm protein identity. From HPLC runs, proteins were quantified by their area under the curve using the calculated extinction coefficient of 2980 M^−1^cm^−1^ for His_6_-tagged proteins and 1490 M^−1^cm^−1^ for proteins without His_6_-tags.

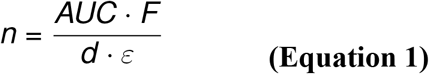

where *n* is the amount of substance of the analyte (in mol), AUC is the area under the curve (in s), *F* is the flow rate (in L/s), *d* is the optical pathlength (in cm), and *ε* is the extinction coefficient (in M^−1^ cm^−1^).

### Steady-state kinetics

Substrate concentrations were chosen to span ≈ 10-fold below and above *K_M_*, as far as not limited by substrate solubility. Optimal starting enzyme concentration (E_0_) and substrate concentration ranges were determined for each variant and substrate combination by empirical sampling. Substrates were pre-dissolved in DMSO at stocks of 200-fold the final concentration, in order to ensure constant DMSO concentration (0.5%) across all substrate concentrations. Upon measurement, aliquots of these substrate stocks were diluted 1:100 in assay buffer (50 mM HEPES-NaOH, 150 mM NaCl, pH 8.0, 5 mM DTT or 1 mM TCEP), of which 100 μL were subsequently mixed with 100 μL of 2-fold concentrated enzyme solution in microtiter plate wells. For substrates with a *p*-nitrophenol leaving group, the progress of the reaction was monitored by absorbance at a wavelength of 405 nm in a spectrophotometric microplate reader (Tecan Infinite 200PRO; Tecan, Switzerland) at 25 °C. The initial rates were extracted by linear fit of the first measurements (at < 10% progress of the reaction) for each substrate concentration and normalised with an extinction coefficient determined from a calibration curve. In order to determine the Michaelis-Menten parameters *k_cat_* and *K_M_*, the data was fitted to the following equation using the non-linear fitting function nls() in R^60^:

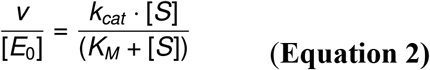

where *v* is the initial rate of the reaction (in mol/s), [*E_0_*] is the initial enzyme concentration (in M), *k_cat_* is the turnover rate (in s^−1^), *K_M_* is the Michaelis constant (in M), and [*S*] is the substrate concentration (in M).

For cyclic nucleotide hydrolysis, unless otherwise indicated, assays used fully reduced proteins in TBS and 1 mM TCEP as described above. For each assay protein aliquots were thawed from storage at −80 °C, diluted to the appropriate concentration, after which metals and substrate were added. Metals and substrate were added from 10X stocks. Nucleotides and metals were in TBS and bis-*p*NPP was in DMSO. All assays included side-by-side samples with substrate, metal, and buffer but no protein to measure autohydrolysis, which was used to baseline the measurements.

Cyclic nucleotide hydrolysis was quantified by UV absorption at 254 nm for AMP and cAMP separated by HPLC, or 260 nm for GMP and cGMP. Assays were performed on an Agilent 1100 series HPLC with a reverse phase column (Agilent Zorbax 300SB-C18, 5 μM, 2.1 × 150 mm). Solvent A was water containing 0.1% TFA and solvent B was acetonitrile containing 0.1% TFA. Elution was isocratic with 3% column B for 5 min, followed by a 5-min flush with solvent B, and a 5-min re-equilibration with 3% solvent B. An example separation is shown in **Figure S9a**. Time-resolved kinetics were followed by 10 μL sample injections using the autosampler module with a 15-min runtime and an isopropanol wash of the needle between injections

Turnover was quantified by the area under the curve for AMP or GMP. An external standard curve **Figure S9b** matched the theoretical signal calculated by **Equation 1**, with the NTP extinction coefficient taken from Cavaluzzi *et al*.^61^. Bis-*p*NPP hydrolysis was measured in microtiter plates using the absorbance at 405 nm measured using a Thermo Scientific Varioskan. Turnover was quantified by an external standard curve using *para*-nitrophenol. dA-P-dA hydrolysis was monitored like the cyclic nucleotides, with turnover calculated using the appearance of both products and the absorbance of the adenine nucleobase present in both products (dA-P and dA). Sample raw data is shown in **Figure S10**.

### Circular Dichroism

CD data were collected on a Chirascan CD spectrometer (Applied Photophysics) from 200 to 260 nm in triplicate and averaged. Far-UV CD spectra were collected using a 1-mm pathlength cuvette and 40 μM mini-cAMPase (or alanine mutant) in Tris/HCl Buffer, or 20 μM S-824 in Tris/HCl Buffer.

### NMR

Protein was concentrated with centrifugal filter units (Amicon, 3 kDa MWCO) to a final concentration of 1 mg/mL in phosphate buffered saline. Proton spectra were collected at 25 °C by using a Bruker Avance III 800 MHz spectrometer. The ^1^H chemical shift was referenced to the DOH line.

### Mutagenesis

PCR mutagenesis was performed by whole plasmid PCR with Q5 high-fidelity polymerase (New England Biolabs) followed by PCR purification (Qiagen) and treatment with KLD Enzyme Mix (New England Biolabs). The Primers were made by Sigma-Aldrich and are listed in **Table S4**. His_6_-tagged proteins included a TEV protease cleavage recognition site.

## ASSOCIATED CONTENT

Additional experimental procedures, figures, and tables can be found in the Supplementary Information. Correspondence and requests for material should be addressed to M.H.H. or F.H.

## AUTHOR INFORMATION

### Corresponding Authors

**Michael H. Hecht**, Department of Chemistry, Princeton University, New Jersey 08540, USA

**Florian Hollfelder**, Department of Biochemistry, University of Cambridge, 80 Tennis Court Road, Cambridge, CB2 1GA, United Kingdom

### Authors

**J. David Schnettler**, Department of Biochemistry, University of Cambridge, 80 Tennis Court Road, Cambridge, CB2 1GA, United Kingdom

*Current address*: Institute of Integrative Biology, Department of Environmental Systems Science, ETH Zurich, Universitätsstrasse 16, 8092 Zürich, Switzerland

**Michael S. Wang**, Department of Chemistry, Princeton University, New Jersey 08540, USA

*Current address*: ZymoChem, 1933 Davis Street, San Leandro, California 94577, USA

**Maximilian Gantz**, Department of Biochemistry, University of Cambridge, 80 Tennis Court Road, Cambridge, CB2 1GA, United Kingdom

**Christina Karas**, Department of Molecular Biology, Princeton University, New Jersey 08540, USA

*Current address*: Modern Meadow, 111 Ideation Way, Suite 100, Nutley, New Jersey 07110, USA

## Acknowledgements

We thank Adrian Bunzel (ETH Zurich) for valuable feedback on the manuscript.

## Author contributions

M.H.H. and J.D.S. initiated the project. C.K. constructed the library. J.D.S. carried out the microfluidic screening. M.S.W. performed the molecular characterization of the hit sequences, with J.D.S. contributing. M.G. performed next-generation sequencing and data analysis. M.H.H. and F.H. directed the work. All authors contributed to the design of the project and discussion of the results. M.S.W., J.D.S., M.G., F.H., and M.H.H. wrote the paper.

## EXTENDED DATA

**Extended Data Figure 1:**
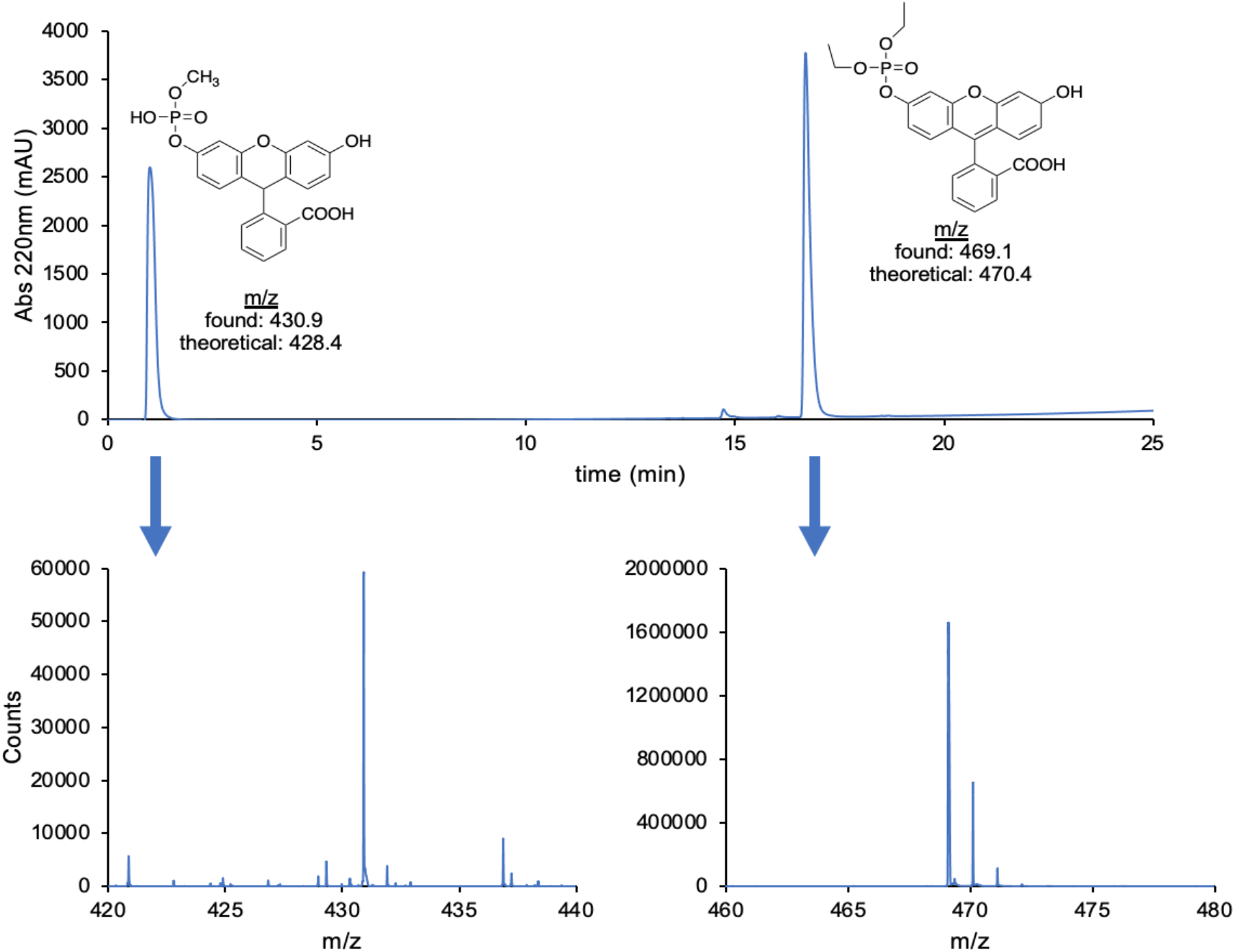
**Fluorogenic substrate mixture** analyzed by LC-MS. The top panel shows the reverse-phase HPLC chromatogram with an increasing gradient of acetonitrile. Below are the positive-mode ESI-MS spectra for each peak, emphasizing the identified mass.

**Extended Data Figure 2:**
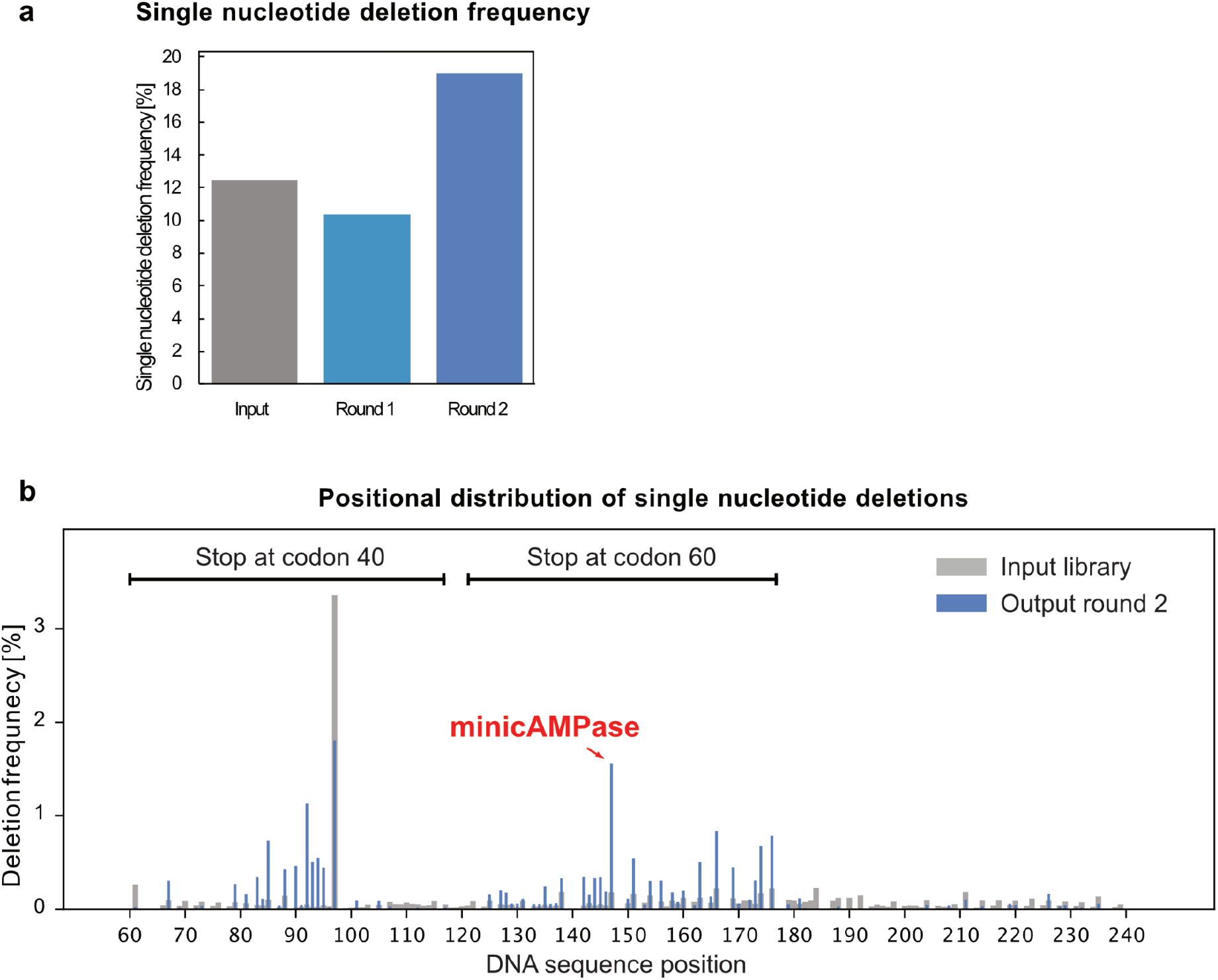
Enrichment of single-nucleotide deletions causing frameshifts after sorting 2. (**a**) Relative frequency (percentage of sequences including one single nucleotide deletion of total number of sequences) of single nucleotide deletions in input library (grey, 12%), after sorting 1 (light blue, 10%) and after sorting 2 (dark blue, 19%) reveals 1.6-fold enrichment of single nucleotide deletions after sorting 2 (**b**) Frequency of single nucleotide deletions at every sequenced position in input library (grey, broad bars) and after sorting 2 (blue, narrow bars). Deletions between position 55 and 117 cause a stop codon at codon 40 while deletions between position 121 and 177 cause a stop codon at codon 60. Deletions at position 147 (as found in the variant mini-cAMPase) are enriched 8.7-fold (0.18 vs 1.56%).

**Extended Data Figure 3:**
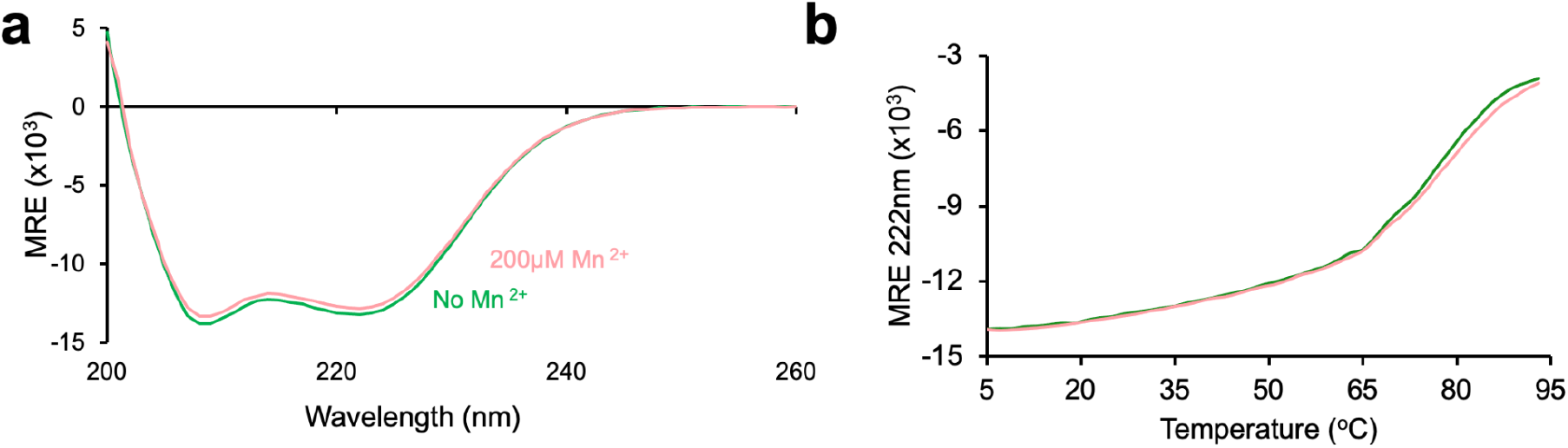
Metal binding effects on structure and thermal stability by Circular Dichroism (CD). (**a**) Full CD spectra of 40 μM mini-cAMPase in TBS without added Mn^2+^ (green) and with added 200 μM Mn^2+^ (pink). (**b**) Melting curves with and without manganese following the same samples in (**a**).

**Extended Data Figure 4:**
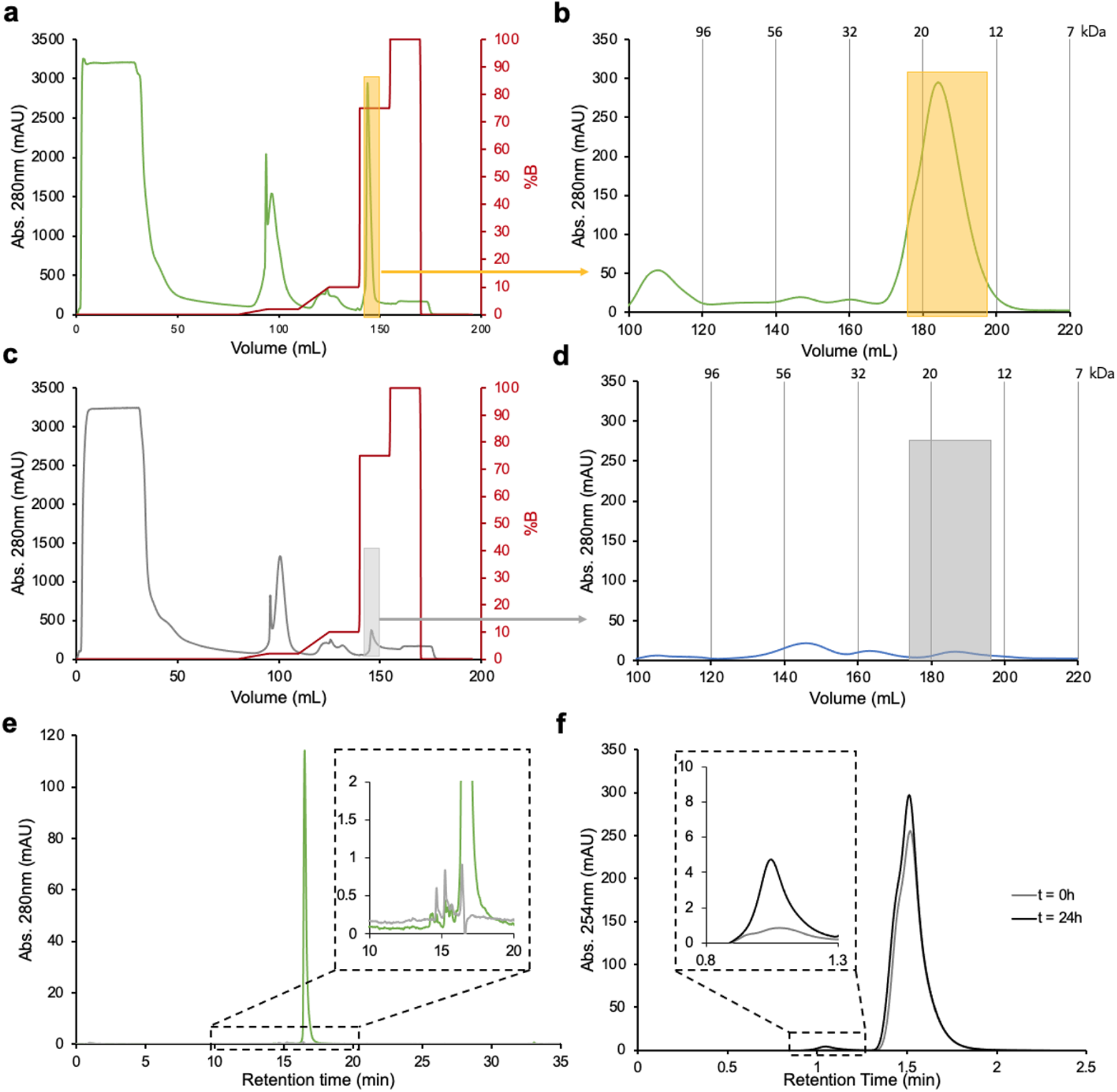
Controls for purification of mini-cAMPase. (**a**) Purification of (His)_6_-tagged mini-cAMPase on the nickel-NTA column, with the taken fractions boxed in yellow. These are then put on (**b**) the preparation-scale sizing column, where the fractions containing the protein are taken (again in yellow). To control for any endogenous proteins that come from this purification steps, in the same cellular background, the un-tagged protein was expressed alongside with the same fractions taken from the nickel-NTA column (**c**) and sizing column (**d**) here shown as grey boxes. (**e**) The samples with (green) and without (grey) the (His)_6_-tag are compared by HPLC, with the main peak being the protein mini-cAMPase. Inset is the magnified baseline. (**f**) The raw data for a single data point of the purification background control in **Figure 4** (Inset is the appearance of AMP): 250 μM cAMP was incubated with 200 μM Mn^2+^ in the background solution – this plot lacks concentration units but is diluted only slightly by the addition of cAMP and Mn^2+^. Note that the dilution of protein samples to 50 μM for the assay means that any contaminants are more concentrated in this sample (with purification yielding 100–250 μM protein, this can be a significant dilution). These are the same conditions used for cAMPase activity characterization as in **Extended Data Figure 6**. The very low observed activity helps rule out an endogenous contaminant.

**Extended Data Figure 5:**
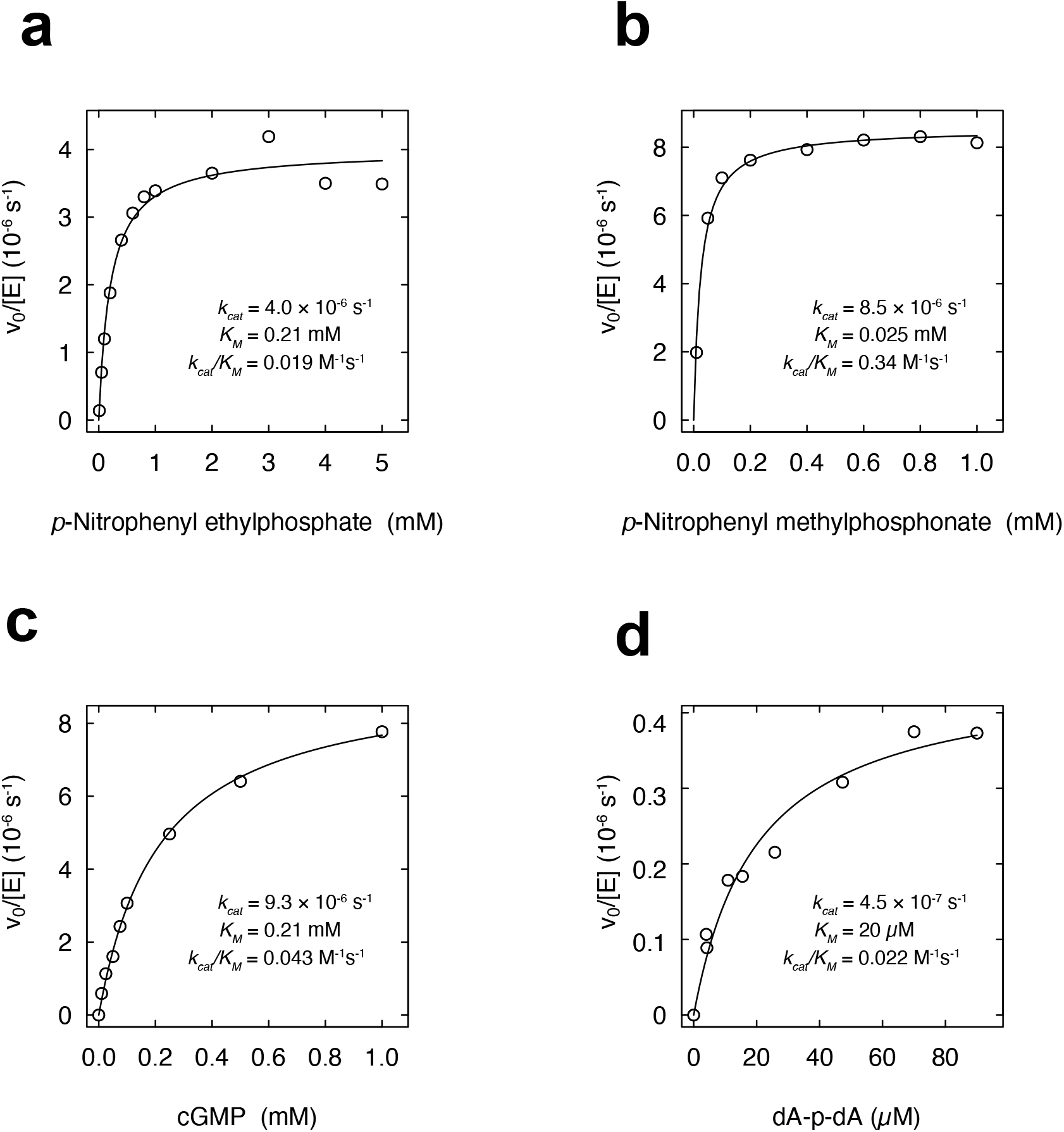
**Michaelis-Menten plots of steady-state kinetics of mini-cAMPase** with **(a)** *p*-nitrophenyl ethylphosphate **(b)** *p*-nitrophenyl methylphosphonate, **(c)** cGMP, and **(d)** dA-P-dA. Protein was lyophilized after HPLC, and resuspended in 50 mM HEPES-NaOH, 150 mM NaCl, 5 mM DTT, pH 8.0 at 25 °C. Kinetics were measured at 50 μM enzyme concentration (25μM dimer) and are calculated per dimer. Curves with cAMP and bis-pNPP are shown in **Figure 4b,c**.

**Extended Data Figure 6:**
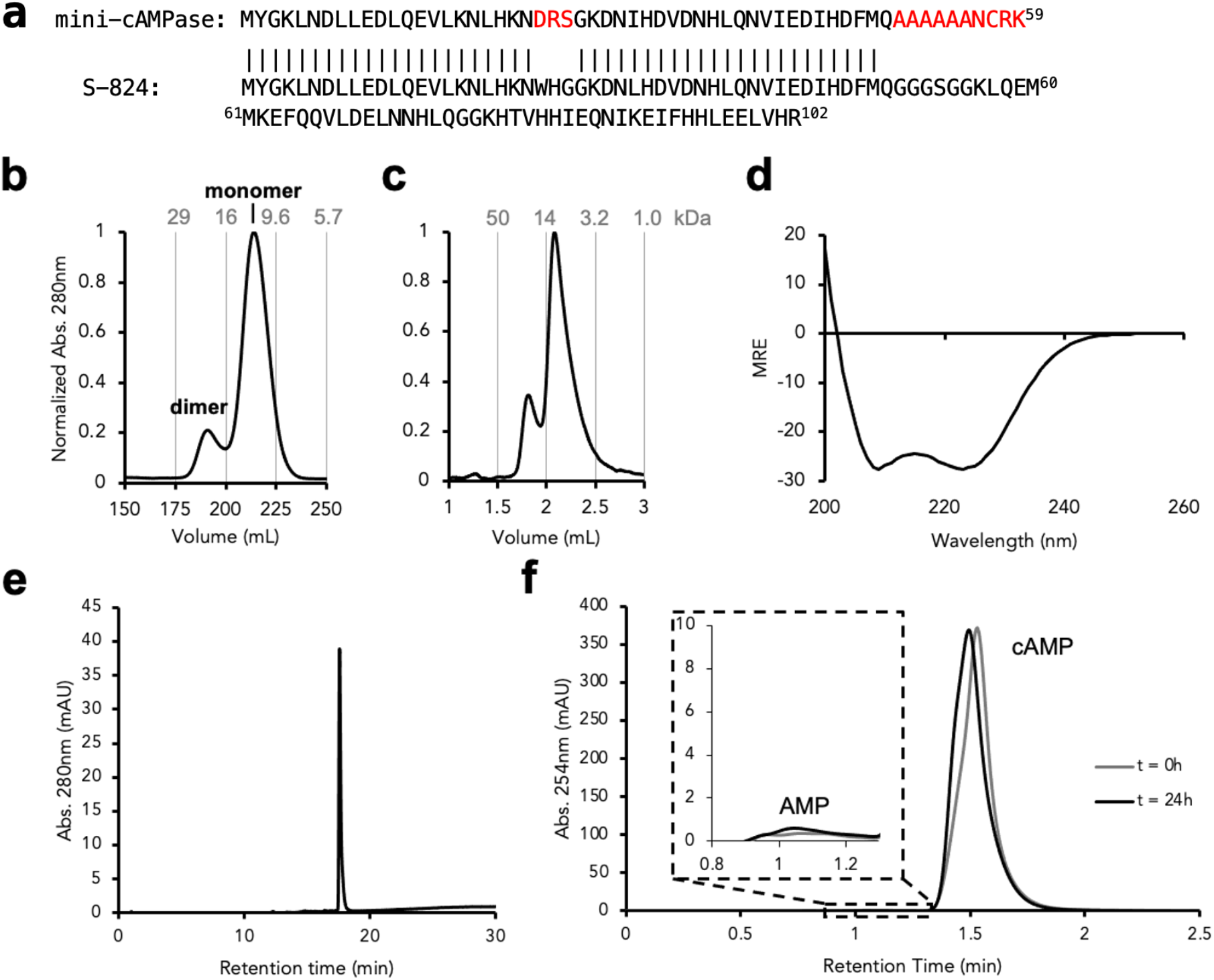
S-824 purification and lack of activity. (**a**) Sequence comparison of characterized hit protein and S-824 with mismatches highlighted. (**b**) Preparation-scale and (**c**) analytical-scale size-exclusion chromatography shows S-824 exists mostly as a monomer, but is partially a dimer. (**d**) Circular Dichroism spectra of S-824 in TBS shows it is helical. (**e**) Reverse-phase HPLC of S-824 shows it is of high purity (> 99%). (**f**) The raw data for a single data point in **Extended Data Figure 8**, sampled before and after 24 h for 50 μM S-824 incubated with 200 μM Mn^2+^ and 250 μM cAMP (the same conditions used for cAMPase activity characterization in **Extended Data Figure 8**), showing that S-824 lacks cAMPase activity.

**Extended Data Figure 7:**
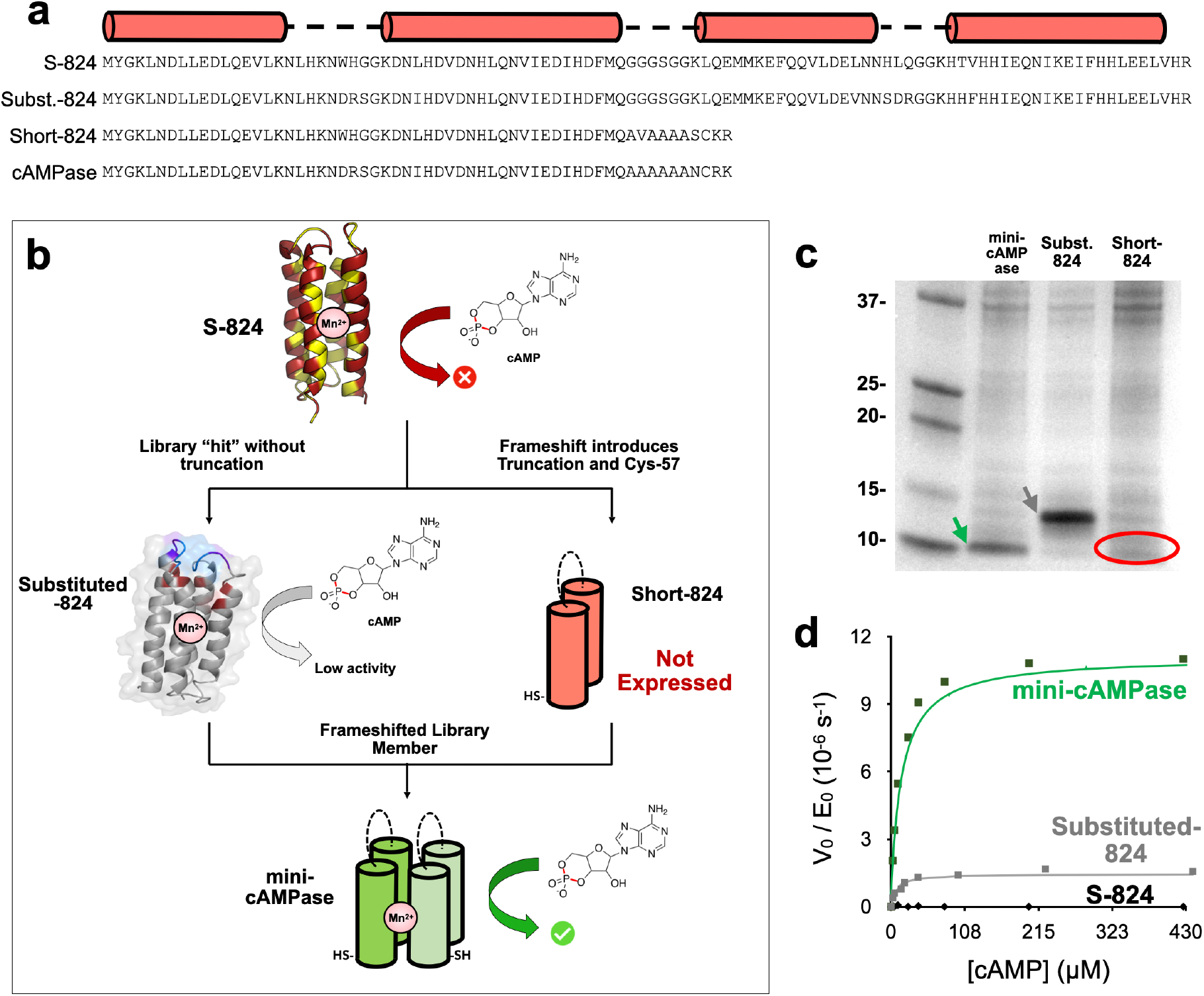
Substitutions and truncations contribute to phosphodiesterase activity. (**a**) Protein sequences showing the changes that bridge S-824 and mini-cAMPase. (**b**) Scheme of the relationship between the sequences and the effect of the truncation on function. (**c**) SDS-PAGE of whole cell extracts shows that Short-824 is poorly expressed. (**d**) Michaelis Menten kinetics comparing S-824, mini-cAMPase, and Substituted-824, showing that Substituted-824 is ≈ 6-fold less active (in *k_cat_*/*K_M_*) than the truncated mini-cAMPase protein.

**Extended Data Table 1:** **Michaelis-Menten kinetics for alanine scanning mutants** measured at 50 μM enzyme concentration in 50 mM HEPES-NaOH, 150 mM NaCl, 200 μM MnCl_2_, pH 8.0 at 25 °C. Note that for truncated proteins, which form dimers, the kinetics are calculated per protein dimer.

| **Protein** | **cAMP** |  |  | **Bis-*p*NPP** |  |  |
| --- | --- | --- | --- | --- | --- | --- |
| | $k_{cat}$<br>($10^{-6} \text{ s}^{-1}$) | $K_M$<br>($\mu\text{M}$) | $k_{cat}/K_M$<br>($\text{M}^{-1}\text{s}^{-1}$) | $k_{cat}$<br>($10^{-6} \text{ s}^{-1}$) | $K_M$<br>($\mu\text{M}$) | $k_{cat}/K_M$<br>($\text{M}^{-1}\text{s}^{-1}$) |
| No Mutation | 22 | 10 | 2.2 | 50 | 160 | 0.3 |
| H20A | 6.2 | 6.5 | 1 | 44 | 440 | 0.1 |
| H31A | 6.2 | 4.4 | 1.4 | 34 | 850 | 0.04 |
| H36A | 7.8 | 6.1 | 1.2 | 44 | 550 | 0.08 |
| H45A | 5.6 | 6.7 | 0.8 | 34 | 650 | 0.06 |
| C57A | 3.4 | 4.6 | 0.8 | 13.4 | 250 | 0.04 |
| R58A | 16 | 10 | 1.6 | 40 | 320 | 0.12 |
| Substituted-824 | 3.2 | 7 | 0.4 | 3 | 235 | 0.01 |
| S-824 | None Detected |  |  | 1.9 | 500 | 0.004 |

**Extended Data Figure 8:**
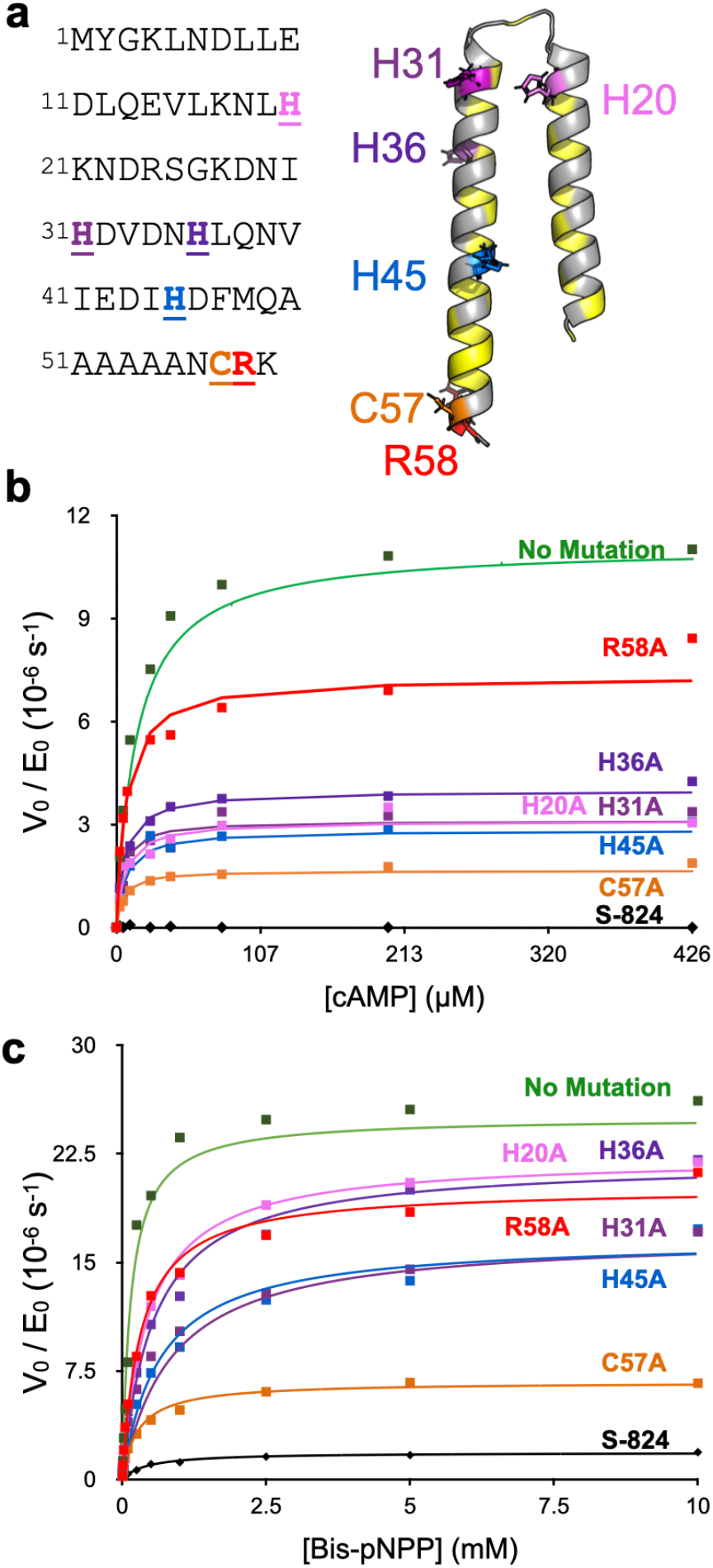
Impact of alanine mutations on the activity of mini-cAMPase. **(a**) Protein sequence and predicted structure with hydrophobic residues in yellow. Sites of mutated residues are emphasized and color coded. (**b**) cAMPase kinetics of alanine point mutants targeting the potential metal binding residues, alongside the inactive ancestor S-824 (black diamonds along baseline, with example raw data in **Ext. Data Fig. 6**). (**c**) Bis-*p*NPP kinetics of alanine point mutants, alongside the inactive ancestor S-824. Note that panels (b) and (c) are plotted on different scales.

**Extended Data Figure 9:**
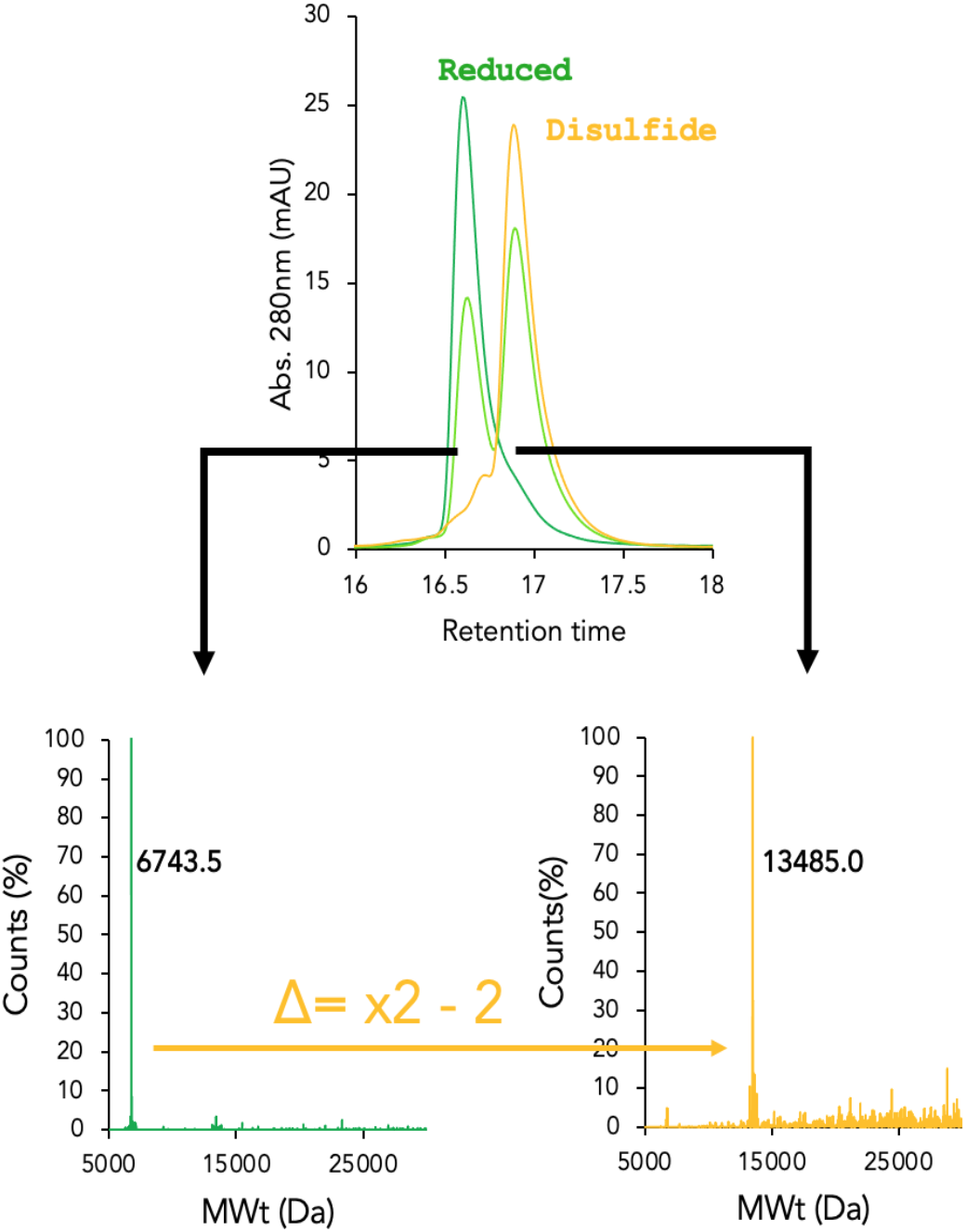
LC-MS of oxidized and reduced protein. Protein peaks were collected after HPLC, analyzed by ESI-MS, and deconvoluted. The deconvoluted spectra correspond to the molecular weight of mini-cAMPase and a disulfide-bonded dimer of mini-cAMPase

## REFERENCES

1. Tiessen, A., Pérez-Rodríguez, P. & Delaye-Arredondo, L. J. Mathematical modeling and comparison of protein size distribution in different plant, animal, fungal and microbial species reveals a negative correlation between protein size and protein number, thus providing insight into the evolution of proteomes. BMC Res. Notes 5, 85 (2012).

2. Mandecki, W. A method for construction of long randomized open reading frames and polypeptides. Protein Eng. 3, 221–226 (1990).

3. Prijambada, I. D. et al. Solubility of artificial proteins with random sequences. FEBS Lett. 382, 21–25 (1996).

4. Keefe, A. D. & Szostak, J. W. Functional proteins from a random-sequence library. Nature 410, 715–718 (2001).

5. Jensen, R. A. Enzyme Recruitment in Evolution of New Function. Annu. Rev. Microbiol. 30, 409–425 (1976).

6. O’Brien, P. J. & Herschlag, D. Catalytic promiscuity and the evolution of new enzymatic activities. Chem. Biol. 6, R91–R105 (1999).

7. Colin, P.-Y. et al. Ultrahigh-throughput discovery of promiscuous enzymes by picodroplet functional metagenomics. Nat. Commun. 6, 10008 (2015).

8. Seelig, B. & Szostak, J. W. Selection and evolution of enzymes from a partially randomized non-catalytic scaffold. Nature 448, 828–831 (2007).

9. Chao, F.-A. et al. Structure and dynamics of a primordial catalytic fold generated by in vitro evolution. Nat. Chem. Biol. 9, 81–83 (2013).

10. Hilvert, D. Design of Protein Catalysts. Annu. Rev. Biochem. 82, 447–470 (2013).

11. Axe, D. D. Estimating the Prevalence of Protein Sequences Adopting Functional Enzyme Folds. J. Mol. Biol. 341, 1295–1315 (2004).

12. Eck, R. V. & Dayhoff, M. O. Evolution of the Structure of Ferredoxin Based on Living Relics of Primitive Amino Acid Sequences. Science 152, 363–366 (1966).

13. Romero Romero, M. L., Rabin, A. & Tawfik, D. S. Functional Proteins from Short Peptides: Dayhoff’s Hypothesis Turns 50. Angew. Chem. Int. Ed. 55, 15966–15971 (2016).

14. Wei, Y., Kim, S., Fela, D., Baum, J. & Hecht, M. H. Solution structure of a de novo protein from a designed combinatorial library. Proc. Natl. Acad. Sci. 100, 13270–13273 (2003).

15. Wei, Y. et al. Stably folded de novo proteins from a designed combinatorial library. Protein Sci. 12, 92–102 (2003).

16. Karas, C. & Hecht, M. A Strategy for Combinatorial Cavity Design in De Novo Proteins. Life 10, 9 (2020).

17. Kamtekar, S., Schiffer, J. M., Xiong, H., Babik, J. M. & Hecht, M. H. Protein design by binary patterning of polar and nonpolar amino acids. Science 262, 1680–1685 (1993).

18. Colin, P.-Y., Zinchenko, A. & Hollfelder, F. Enzyme engineering in biomimetic compartments. Curr. Opin. Struct. Biol. 33, 42–51 (2015).

19. Gantz, M., Aleku, G. A. & Hollfelder, F. Ultrahigh-throughput screening in microfluidic droplets: a faster route to new enzymes. Trends Biochem. Sci. S096800042100236X (2021) doi:10.1016/j.tibs.2021.11.001.

20. Baret, J.-C. et al. Fluorescence-activated droplet sorting (FADS): efficient microfluidic cell sorting based on enzymatic activity. Lab. Chip 9, 1850 (2009).

21. Jumper, J. et al. Highly accurate protein structure prediction with AlphaFold. Nature 596, 583–589 (2021).

22. Mirdita, M. et al. ColabFold: making protein folding accessible to all. Nat. Methods 19, 679–682 (2022).

23. Fowler, D. M. et al. High-resolution mapping of protein sequence-function relationships. Nat. Methods 7, 741–746 (2010).

24. Hietpas, R. T., Jensen, J. D. & Bolon, D. N. A. Experimental illumination of a fitness landscape. Proc. Natl. Acad. Sci. 108, 7896–7901 (2011).

25. Larsen, A. C. et al. A general strategy for expanding polymerase function by droplet microfluidics. Nat. Commun. 7, 11235 (2016).

26. Check Hayden, E. Chemistry: Designer debacle. Nature 453, 275–278 (2008).

27. O’Brien, P. J. & Herschlag, D. Functional Interrelationships in the Alkaline Phosphatase Superfamily: Phosphodiesterase Activity of *Escherichia coli* Alkaline Phosphatase. Biochemistry 40, 5691–5699 (2001).

28. Imamura, R. et al. Identification of the cpdA Gene Encoding Cyclic 3ʹ,5ʹ-Adenosine Monophosphate Phosphodiesterase in Escherichia coli. J. Biol. Chem. 271, 25423–25429 (1996).

29. van Loo, B. et al. An efficient, multiply promiscuous hydrolase in the alkaline phosphatase superfamily. Proc. Natl. Acad. Sci. 107, 2740–2745 (2010).

30. Schroeder, G. K., Lad, C., Wyman, P., Williams, N. H. & Wolfenden, R. The time required for water attack at the phosphorus atom of simple phosphodiesters and of DNA. Proc. Natl. Acad. Sci. 103, 4052–4055 (2006).

31. Chin, J. & Zou, X. Catalytic hydrolysis of cAMP. Can. J. Chem. 65, 1882–1884 (1987).

32. Bar-Even, A. et al. The Moderately Efficient Enzyme: Evolutionary and Physicochemical Trends Shaping Enzyme Parameters. Biochemistry 50, 4402–4410 (2011).

33. Bar-Even, A., Milo, R., Noor, E. & Tawfik, D. S. The Moderately Efficient Enzyme: Futile Encounters and Enzyme Floppiness. Biochemistry 54, 4969–4977 (2015).

34. Copley, S. D., Newton, M. S. & Widney, K. A. How to Recruit a Promiscuous Enzyme to Serve a New Function. Biochemistry (2022) doi:10.1021/acs.biochem.2c00249.

35. Radzicka, A. & Wolfenden, R. A proficient enzyme. Science 267, 90–93 (1995).

36. Chen, J. et al. An Asymmetric Dizinc Phosphodiesterase Model with Phenolate and Carboxylate Bridges. Inorg. Chem. 44, 3422–3430 (2005).

37. Matange, N. Revisiting bacterial cyclic nucleotide phosphodiesterases: cyclic AMP hydrolysis and beyond. FEMS Microbiol. Lett. 362, fnv183 (2015).

38. Studer, S. et al. Evolution of a highly active and enantiospecific metalloenzyme from short peptides. Science 362, 1285–1288 (2018).

39. Basler, S. et al. Efficient Lewis acid catalysis of an abiological reaction in a de novo protein scaffold. Nat. Chem. 1–5 (2021) doi:10.1038/s41557-020-00628-4.

40. Razkin, J., Lindgren, J., Nilsson, H. & Baltzer, L. Enhanced Complexity and Catalytic Efficiency in the Hydrolysis of Phosphate Diesters by Rationally Designed Helix-Loop-Helix Motifs. ChemBioChem 9, 1975–1984 (2008).

41. Seelig, B. mRNA display for the selection and evolution of enzymes from in vitro-translated protein libraries. Nat. Protoc. 6, 540–552 (2011).

42. Ma, B. & Nussinov, R. Enzyme dynamics point to stepwise conformational selection in catalysis. Curr. Opin. Chem. Biol. 14, 652–659 (2010).

43. Nashine, V. C., Hammes-Schiffer, S. & Benkovic, S. J. Coupled motions in enzyme catalysis. Curr. Opin. Chem. Biol. 14, 644–651 (2010).

44. Kern, D. From structure to mechanism: skiing the energy landscape. Nat. Methods 18, 435–436 (2021).

45. Dellus-Gur, E. et al. Negative Epistasis and Evolvability in TEM-1 β-Lactamase—The Thin Line between an Enzyme’s Conformational Freedom and Disorder. J. Mol. Biol. 427, 2396–2409 (2015).

46. Mabbitt, P. D. et al. Conformational Disorganization within the Active Site of a Recently Evolved Organophosphate Hydrolase Limits Its Catalytic Efficiency. Biochemistry 55, 1408–1417 (2016).

47. Tokuriki, N. & Tawfik, D. S. Protein Dynamism and Evolvability. Science 324, 203–207 (2009).

48. Campbell, E. et al. The role of protein dynamics in the evolution of new enzyme function. Nat. Chem. Biol. 12, 944–950 (2016).

49. Vamvaca, K., Vögeli, B., Kast, P., Pervushin, K. & Hilvert, D. An enzymatic molten globule: Efficient coupling of folding and catalysis. Proc. Natl. Acad. Sci. 101, 12860–12864 (2004).

50. Smith, B. A., Mularz, A. E. & Hecht, M. H. Divergent evolution of a bifunctional de novo protein. Protein Sci. Publ. Protein Soc. 24, 246–252 (2015).

51. Bershtein, S., Segal, M., Bekerman, R., Tokuriki, N. & Tawfik, D. S. Robustness–epistasis link shapes the fitness landscape of a randomly drifting protein. Nature 444, 929–932 (2006).

52. Yang, G. et al. Higher-order epistasis shapes the fitness landscape of a xenobiotic-degrading enzyme. Nat. Chem. Biol. 15, 1120–1128 (2019).

53. Kaltenbach, M., Jackson, C. J., Campbell, E. C., Hollfelder, F. & Tokuriki, N. Reverse evolution leads to genotypic incompatibility despite functional and active site convergence. eLife 4, e06492 (2015).

54. Park, Y., Metzger, B. P. H. & Thornton, J. W. Epistatic drift causes gradual decay of predictability in protein evolution. Science 376, 823–830 (2022).

55. Skerra, A. Use of the tetracycline promoter for the tightly regulated production of a murine antibody fragment in Escherichia coli. Gene 151, 131–135 (1994).

56. Schnettler, J. D., Klein, O. J., Kaminski, T. S., Colin, P.-Y. & Hollfelder, F. Ultrahigh-Throughput Directed Evolution of a Metal-Free α/β-Hydrolase with a Cys-His-Asp Triad into an Efficient Phosphotriesterase. J. Am. Chem. Soc. (2022) doi:10.1021/jacs.2c10673.

57. Neun, S., Kaminski, T. S. & Hollfelder, F. Chapter Five - Single-cell activity screening in microfluidic droplets. in Methods in Enzymology (eds. Allbritton, N. L. & Kovarik, M. L.) vol. 628 95–112 (Academic Press, 2019).

58. Faure, A. J., Schmiedel, J. M., Baeza-Centurion, P. & Lehner, B. DiMSum: an error model and pipeline for analyzing deep mutational scanning data and diagnosing common experimental pathologies. Genome Biol. 21, 207 (2020).

59. Rehder, D. S. & Borges, C. R. Cysteine sulfenic Acid as an Intermediate in Disulfide Bond Formation and Nonenzymatic Protein Folding. Biochemistry 49, 7748–7755 (2010).

60. R Core Team. R: A Language and Environment for Statistical Computing. (R Foundation for Statistical Computing, 2017).

61. Cavaluzzi, M. J. & Borer, P. N. Revised UV extinction coefficients for nucleoside-5′-monophosphates and unpaired DNA and RNA. Nucleic Acids Res. 32, e13 (2004).

