## Supplementary Information for "Selection of a Promiscuous Minimalist cAMP Phosphodiesterase from a Library of *De Novo* Designed Proteins"

J. David Schnettler[1,*], Michael S. Wang[2,*], Maximilian Gantz[1], Christina Karas[3], Florian Hollfelder[1, #], Michael H. Hecht[2, #]

[1] Department of Biochemistry, University of Cambridge
[2] Department of Chemistry, Princeton University
[3] Department of Molecular Biology, Princeton University

[*] Both authors contributed equally
[#] Both authors jointly supervised this work

### 1. SUPPLEMENTARY METHODS

#### 1.1 Abbreviations

Adenosine monophosphate (AMP), cyclic adenosine monophosphate (cAMP), circular dichroism (CD), cyclic guanosine monophosphate (cGMP), bis(*p*-nitrophenyl) phosphate (bis-pNPP), *Escherichia coli* (*E. coli*), 4-(2-hydroxyethyl)-1-piperazineethanesulfonic acid (HEPES), isopropyl-β-D-thiogalactoside (IPTG), Miller-Luria Broth (LB), nuclear magnetic resonance (NMR), *para*-nitrophenol (*p*NP), 1H,1H,2H,2H-perfluoro-1-octanol (PFO), terrific broth (TB), melting temperature ($T_M$), trifluoroacetic acid (TFA), and tris(hydroxymethyl) aminomethane (Tris).

#### 1.2 Reagents, Buffers, and Strains

All chemical reagents were purchased from Merck KGaA (Darmstadt, Germany; previously Sigma-Aldrich, St. Louis, MO, USA) and all biological reagents from New England Biolabs (Ipswich, MA, USA), unless otherwise noted. Buffers had the following compositions: HEPES buffer 50 mM HEPES-NaOH, 150 mM NaCl, pH 8.0), Phosphate buffer (50 mM $Na_2HPO_4$, 300 mM NaCl, pH 8.0), Tris buffer (50 mM Tris-NaOH, 300 mM NaCl, pH 8.0). Library screening was carried out in the *E. coli* strain E. cloni 10G. Plasmid preparation and cloning of single hits was carried out in the strain DH5α. Protein expression for purification was carried out in strains BL21(DE3).

#### 1.3 Substrate synthesis

The bait substrate mixture was synthesised by Mark Mohamed as previously described for fluorescein di(diethylphosphate)[1]. Mass spectrometric analysis showed that the reaction product had partly de-alkylated over time, containing proportions of the corresponding phosphodi-, and -triester, so that a mixture of phosphoesters was used for screening (**Ext. Data Fig. 1**).

#### 1.4 Next-Generation Sequencing Analysis: CRX motif analysis

In addition to the enrichment of truncations, we observed that most truncated variants contain a C-terminal Cys-Arg-Xaa (CRX) motif (**Figure S2**). The parent sequence of the library, S-824, is devoid of cysteine, but 15% of all reads in the input library and 22% of reads after sorting 2 contain at least one cysteine (**Figure S2b**). Separating the read counts in truncated and full-length sequences showed that cysteines are only enriched among truncated sequences (2-fold among truncated vs 0.7-fold among full-length sequences). Analysis of position-dependent enrichment of cysteine after sorting 2 revealed enrichment at position 37 and 57 (1.3-fold and 2.8-fold; **Figure S2a**). Additionally, after sorting 2 reads with arginine at position 38 and 58 are enriched 1.3-fold and 2.7-fold, respectively. In total, the C-terminal Cys-Arg-Xaa motif occurs in 17% of reads after sorting 2 which is a 1.7-fold enrichment compared to the input library (10%). In conclusion, the enrichment of frameshifted variants leads to a co-enrichment of truncations and CRX motifs (**Figure S2c**).

#### 1.5 Control experiments: The observed phosphodiesterase activity is not the result of contamination

The observation of catalytic activity for a *de novo* protein expressed in *E. coli* inevitably raises concerns about the possibility of contaminating activity from endogenous proteins[2,3]. To rule out this possibility, we (i) considered the activity of the endogenous *E. coli* cAMPase, and (ii) performed several control experiments. First, consideration of *E. coli* cAMPase (CpdA)[4] suggests differences: CpdA requires $Fe^{2+}$ or $Mg^{2+}$ and has $K_M \approx 500$ µM. In contrast, mini-cAMPase uses an unusual metal cofactor ($Mn^{2+}$), is inactive with $Mg^{2+}$ (**Figure 3a**), and has a lower $K_M \approx 10$ µM.

Nonetheless, we continued to consider the possibility that activity might be due to a contaminating endogenous protein. Therefore, we performed measurements on two biological replicates and showed they had similar activity with both cAMP and the phosphodiesterase substrate bis(*p*-nitrophenyl) phosphate (bis-*p*NPP) (**Figure 3b,c**). We also showed that activity persisted after an additional denaturing purification step using RP-HPLC and lyophilization (Kinetics shown in **Ext. Data Fig. 5** were measured after denaturing purification).

Next, to rule out the possibility that an endogenous *E. coli* protein might co-purify with mini-cAMPase (in biological replicates and in RP-HPLC), we engineered two versions of mini-cAMPase that would purify in completely different fractions. This was accomplished by creating both $His_6$-tagged and untagged versions of mini-cAMPase. In each case, the enzymatic activity co-purified with the *de novo* protein, while the 'dummy' fraction, where the alternate (tagged or untagged) version would have eluted, was inactive (**Figure 3b,c**)[5]. Detailed comparisons of tagged/untagged proteins are shown in **Ext. Data Fig. 4.**

Finally, we demonstrated explicitly that the catalytic activity of mini-cAMPase correlates with the sequence of mini-cAMPase. Thus, the ancestor of the library, S-824, purified in the same way, shows no activity as a cAMP phosphodiesterase. Moreover, mutations causing changes in the sequence of mini-cAMPase produced corresponding changes in activity. Together, these controls and comparisons provide compelling evidence that mini-cAMPase is a *de novo* phosphodiesterase.

#### 1.6 Note on the calculation of rate accelerations and catalytic proficiencies

For the calculation of rate accelerations and catalytic proficiencies for phosphodiester substrates, several uncatalyzed background hydrolysis rates ($k_{uncat}$) are available from the literature (**Table S2**). Chin and Zou[6] estimated the uncatalyzed hydrolysis rate for cAMP as $k_{uncat} \approx 3 \times 10^{-15}$ $s^{-1}$ at pH 7 and 25 °C, extrapolated from the uncatalyzed hydrolysis of the cyclic phosphodiester ethylene phosphate at pH 7 and 100 °C. As this is the only available literature value based on hydrolysis of a cyclic phosphodiester we consider it the most accurate for cAMP. It should be noted, however, that these values do not take into account that hydrolysis of phosphodiesters at high temperatures (at which slow background rates are measured) mostly happens through C–O cleavage. These values therefore merely represent an upper boundary for the rate of P–O cleavage. Schroeder *et al.* re-measured phosphodiester background hydrolysis rates with a sterically hindered substrate where only P–O cleavage can occur (dineopentyl phosphate) and determined a background hydrolysis value of $k_{uncat} \approx 7 \times 10^{-16}$ $s^{-1}$ at pH 7 and 25 °C[7].

#### 2. SUPPLEMENTARY FIGURES

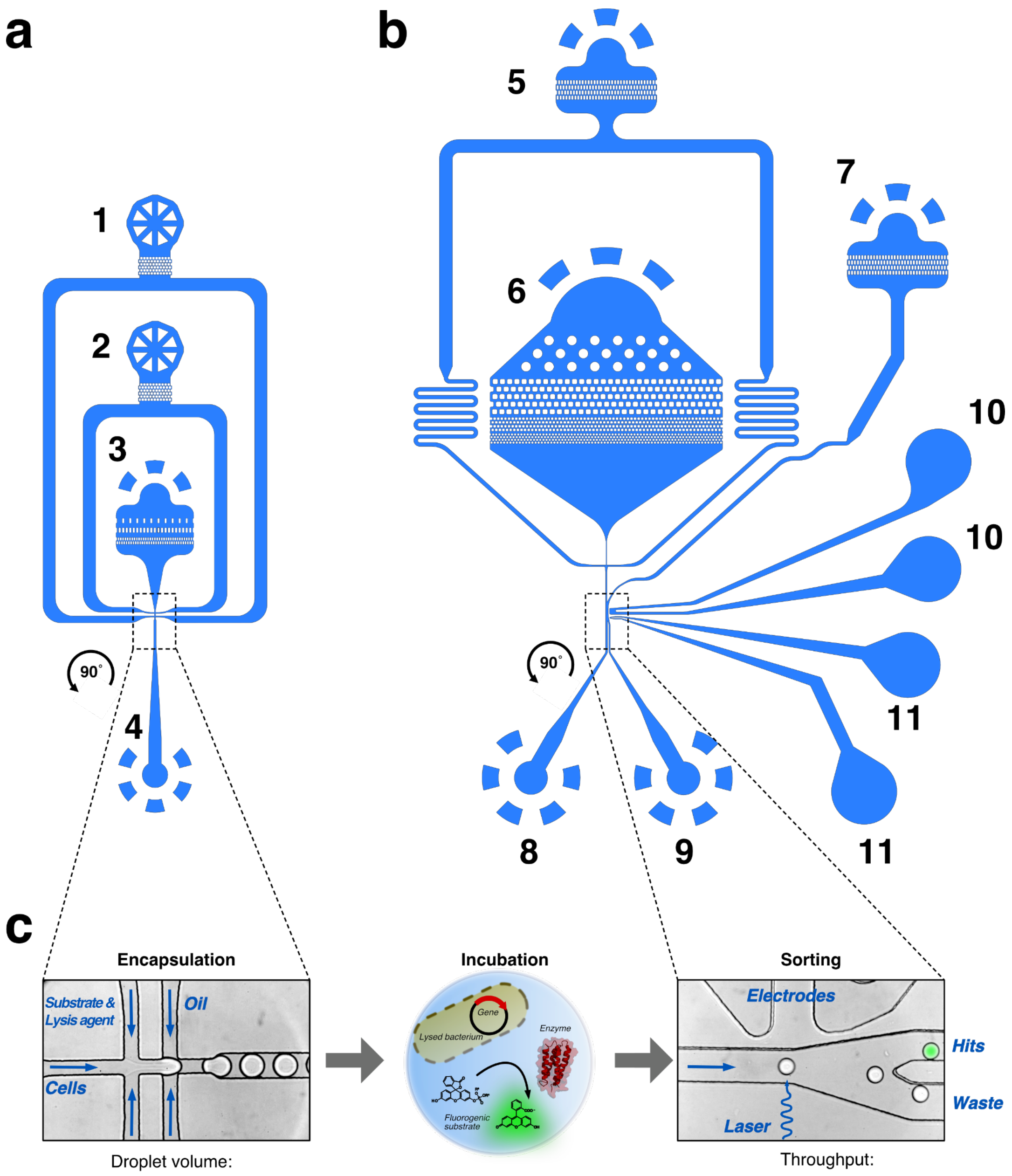


**Figure S1**: **Layouts of microfluidic chips for droplet generation and sorting. (a)** Flow-focussing chip (depth: 12 µm) for droplet generation with (1) oil/surfactant mixture inlet, (2) inlet for substrate/lysis agent mixture, (3) inlet for cell suspension, and (4) outlet for droplet collection. **(b)** Droplet sorting chip (depth: 20 µm) for fluorescence-activated droplet sorting with (5) inlet for spacing oil, (6) inlet for droplets, (7) oil extractor, (8) waste outlet, (9) hit outlet, (10) ground electrode (+), and (11) signal electrode (–). **(c)** On the flow-focusing chip, *E. coli* cells can be co-encapsulated with a fluorogenic substrate and lysis agent into monodisperse picolitre-sized droplets. After incubation, the library-containing emulsion can be screened on the sorting chip, where the droplets pass through a sorting junction with an excitation laser and a fluorescence detector. Upon surpassing a pre-set fluorescence threshold, specific droplets can be electrophoretically sorted into the hit channel for subsequent recovery. Panels **(a)** and **(b)** of this figure are inspired by the work of Neun *et al.*[8].

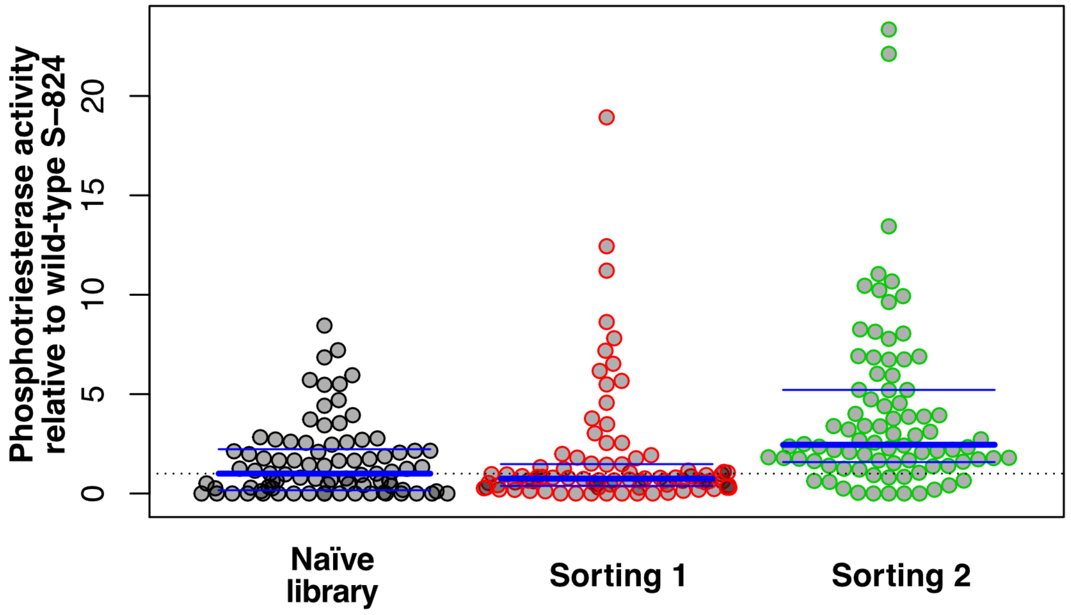


**Figure S2: Secondary screening in microtiter plates shows a cumulative enrichment of clones with phosphoesterase activity**. Lysate activity levels are shown for 84 library clones randomly picked before microfluidic droplet screening and after sorting 1 and sorting 2, respectively. The dotted line indicates the background activity level of wild-type S-824, the thick blue line indicates median activity and the thin blue lines indicate the first and the third quartile.

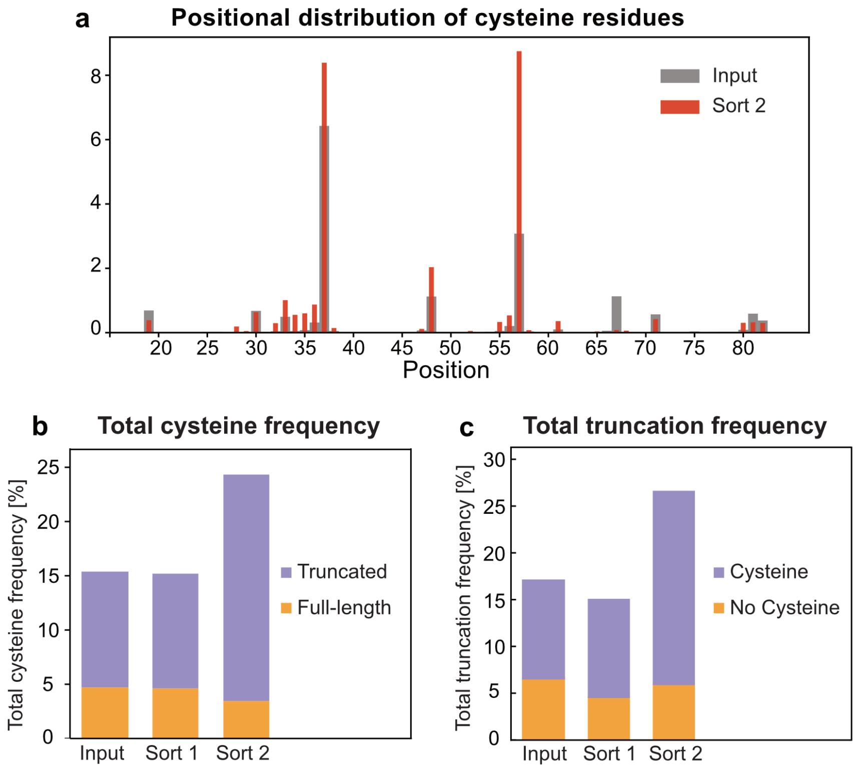


**Figure S3: Occurrence of cysteine residues is enriched at positions 37 and 57 and enrichment is interdependent with truncation**. (a) Frequency of cysteine residues at every sequenced position in the input library (grey, broad bars) and after sorting 2 (red, narrow bars) shows 1.3 and 2.8-fold enrichments of truncations at position 40 and 60 after sorting 2. (b) Relative frequency of reads containing one or more cysteine residues stays constant after sorting 1 (15%) and increases after sorting 2 (15% to 22%). However, the effect is only caused by truncated sequences (violet; 11% vs 18%) while the relative frequency of cysteines even slightly decreases in full-length sequences (orange; 5% vs 4%) (c) The frequency of truncated variants slightly decreases after sorting 1 (17% vs 15%) and increases after sorting 2 (17% vs 27%). However, the enrichment after sorting 2 is only caused by sequences containing cysteine residues (violet, 11% vs 21%), while the fraction of reads *without* cysteine stays constant (orange, 7% vs 6%).

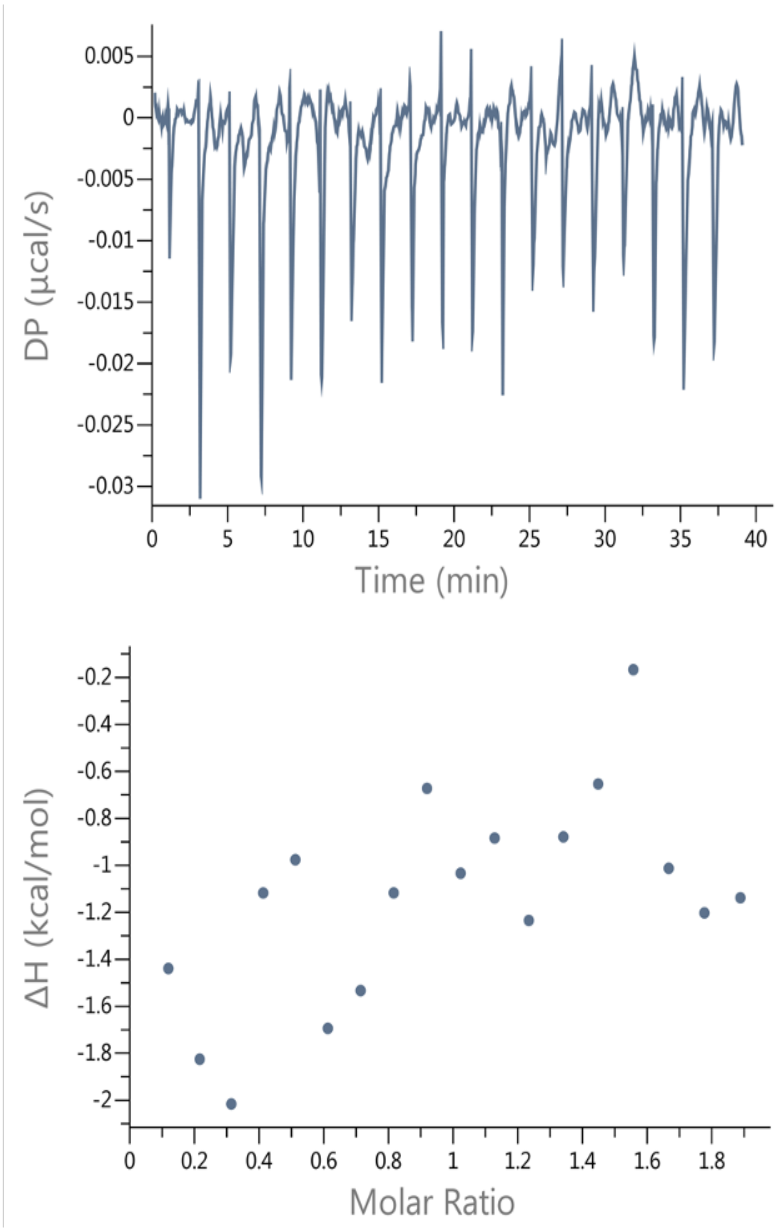


**Figure S4: Metal binding by isothermal titration calorimetry** (ITC) shows no clear binding. Top: raw data, bottom: integrated heat. 10 µM mini-cAMPase was titrated with up to 20 µM $MnCl_2$, but no binding was observed.

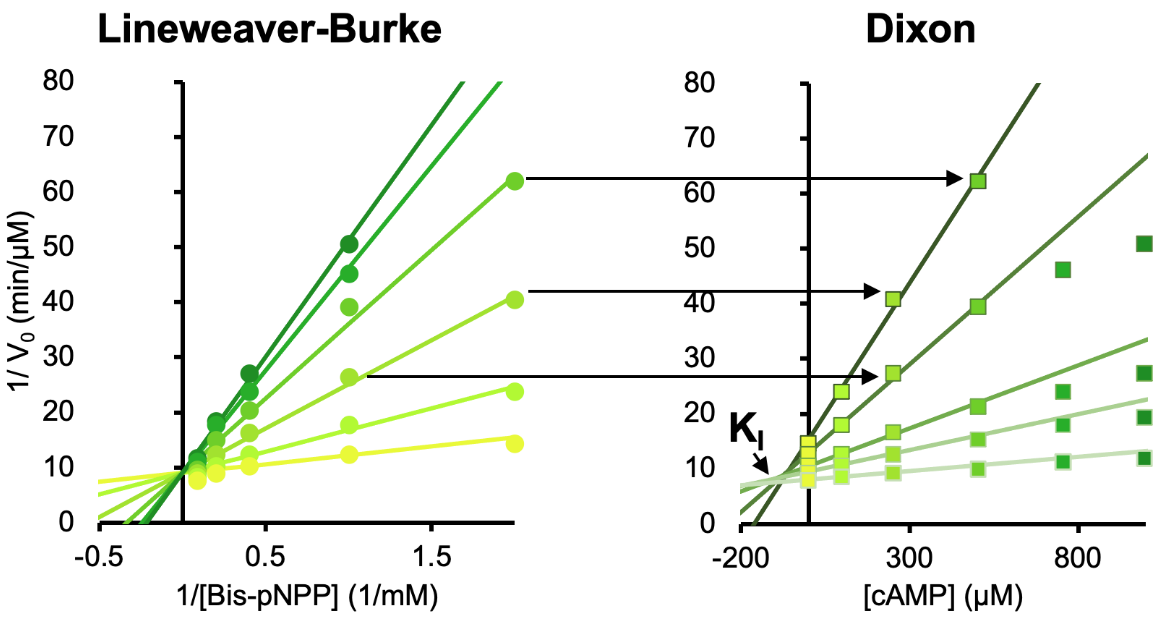


**Figure S5: Inhibition of Bis(PNPP) hydrolysis by cAMP**. On the left is the Lineweaver-Burke data shown in Figure 4d. To the right is this data transformed to a Dixon plot with data colored to match. Lines connect the kinetics at the same concentration of bis(PNPP) substrate, and their intersection point is the $K_i$ of 70 ± 8 $\mu$M cAMP.

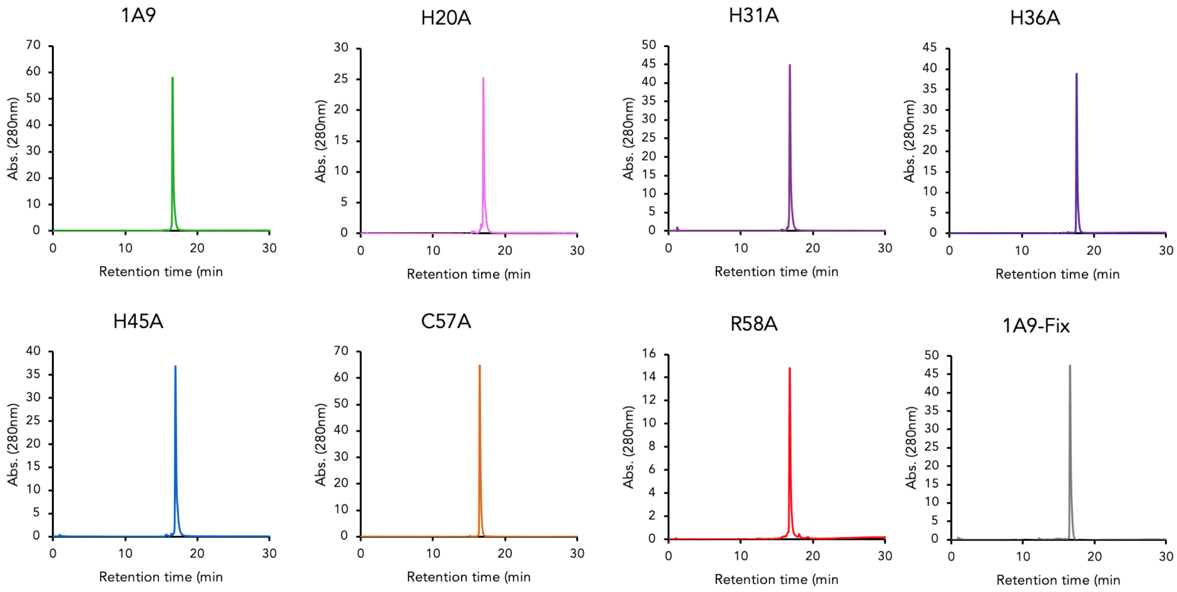


**Figure S6: Protein purity by reverse-phase HPLC**. Reduced alanine point mutants were run as-is after size exclusion on a reverse phase HPLC to separate proteins. Color coding matches in-text figures. In each sample, the main peak is the reduced protein of interest (as confirmed by mass spectrometry).

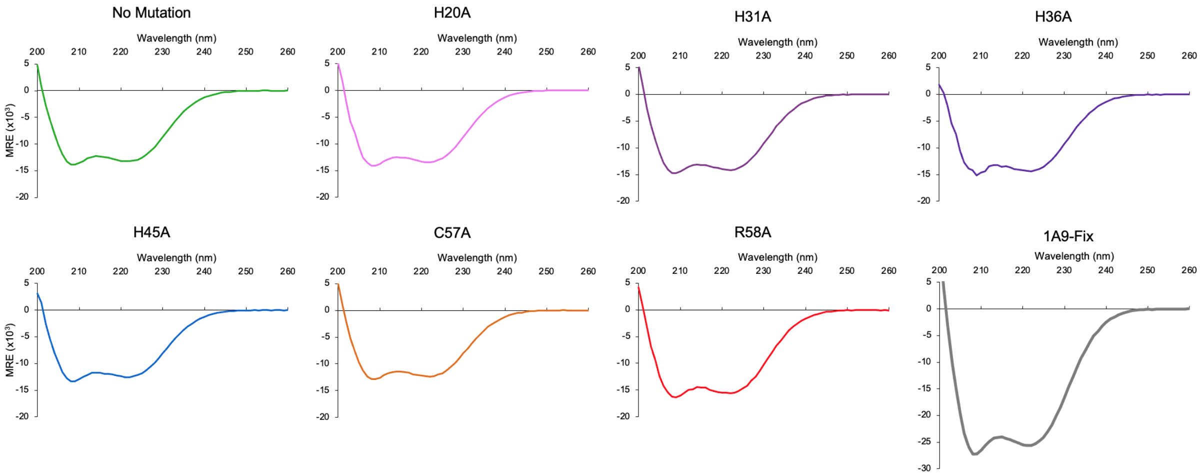


**Figure S7: Effects of mutation on secondary structure** for 40 μM protein. Shown are the CD spectra of the reduced alanine point mutants and mini-cAMPase-Fix. Color coding matches **Extended Data Figure 8** and **Figure S7**. The spectra are comparable to the spectra without mutation, suggesting that the mutations do not significantly alter secondary structure.

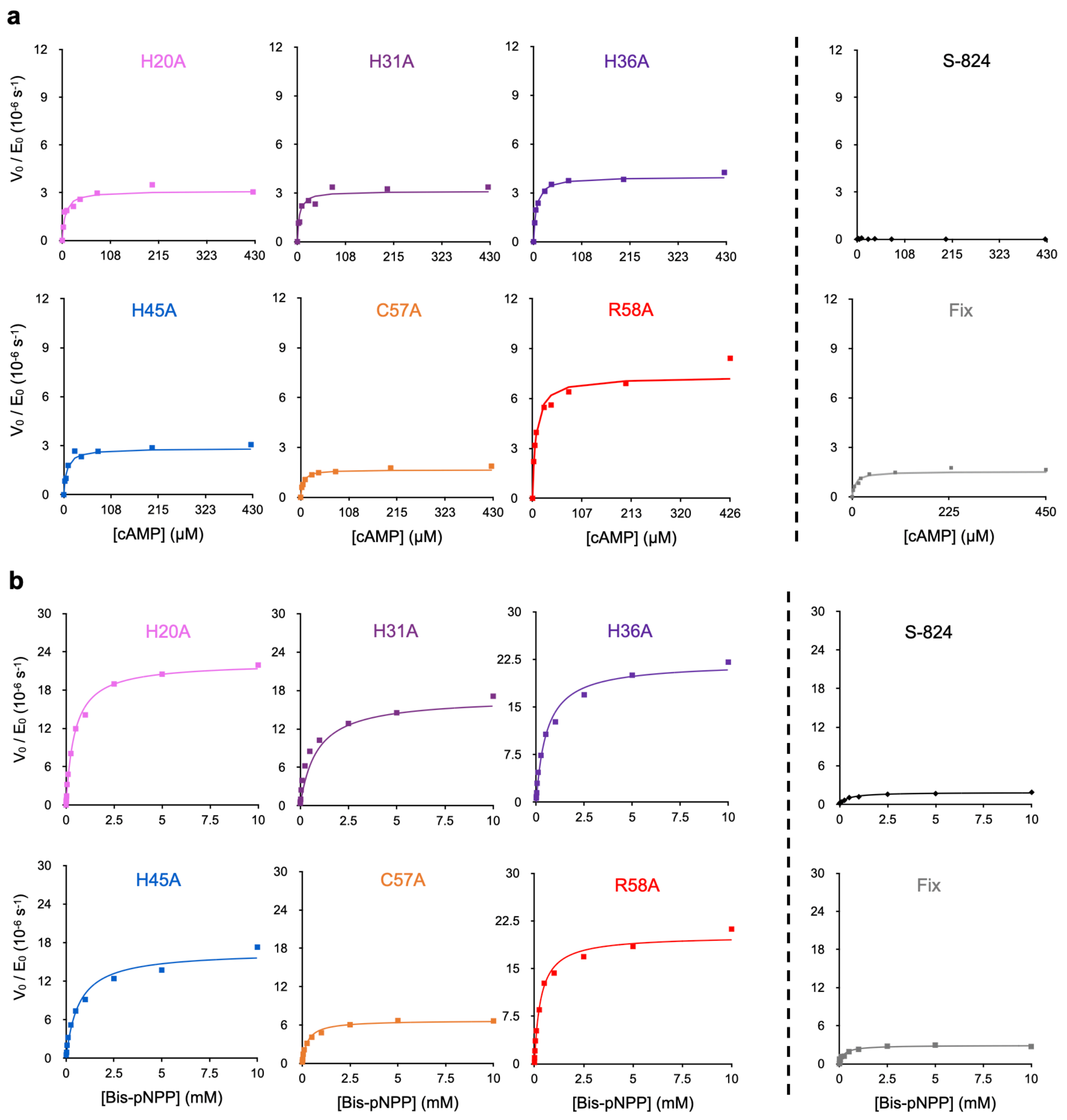


**Figure S8**: **Individual Michaelis-Menten plots** of steady-state kinetics for mutants of mini-cAMPase and S-824. On the left are alanine-scanning mutants, on the right are S-824 and mini-cAMPase-Fix (**a**) with cAMP and (**b**) bis-*p*NPP. These are the same plots as in **Extended Data Figure 8**.

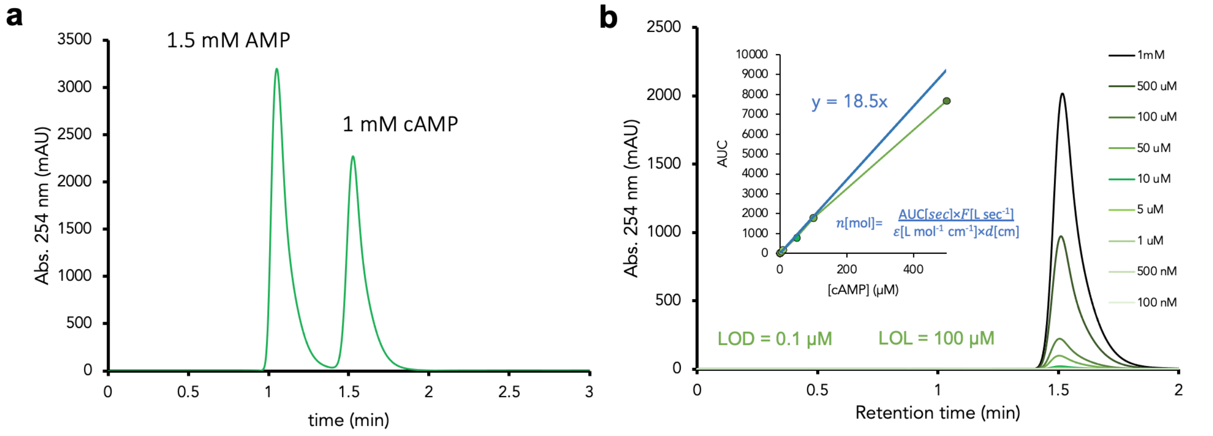


**Figure S9: AMP standards on the HPLC** (**a**) Resolution of AMP (peak 1) and cAMP (peak 2) shows that product and substrate are well resolved even at concentrations above those used in the assays for kinetic characterization. (**b**) External calibration of cAMP concentration, showing the raw data for the cAMP peak with standard curve inset. Variable concentrations of cAMP are quantifiable by the area under the curve. This matches the theoretical calibration curve with known extinction coefficient for cAMP up until 100µM cAMP. This makes the limit of linearity (LOL) 100µM for this analytical method. The limit of detection (3**σ**, LOD) is also the lowest standard concentration of 0.1µM.

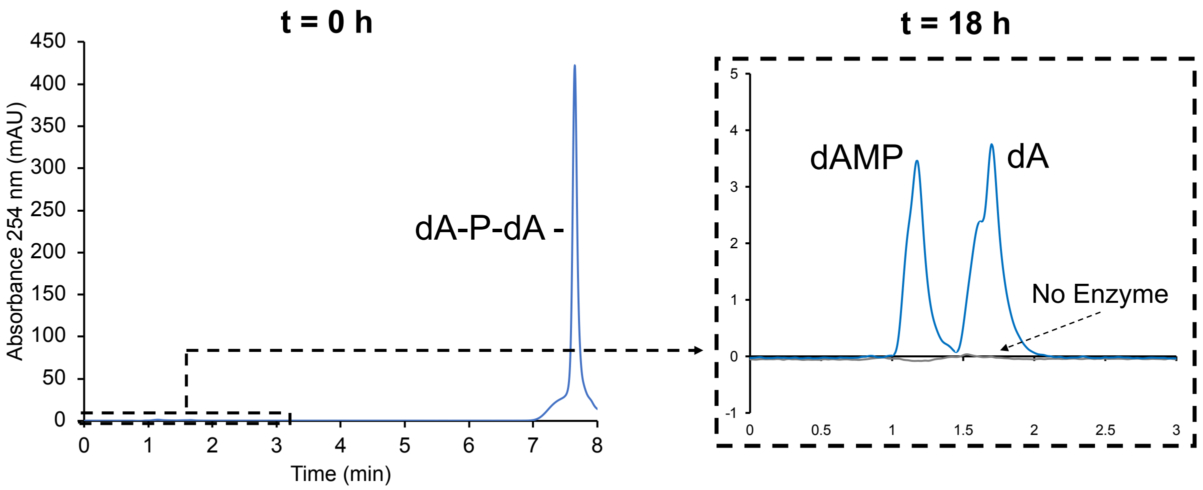


**Figure S10: Raw data for nuclease activity.** 100 µM deoxyadenosine dinucleotide (dA-P-dA) was incubated with 100 µM mini-cAMPase and 200 µM $MnCl_2$ (blue) or TBS buffer and 200 µM $MnCl_2$ (no-enzyme control, grey). Products were analyzed with the same HPLC gradient as other nucleotides. On the left is the RP-HPLC profile after 0 h with the main peak being dA-P-dA. On the right is the magnified product profile at t = 18 h, with the baseline at t = 0 h subtracted, showing slow, enzyme-concentration-dependent turnover.

#### 3. SUPPLEMENTARY TABLES

**Table S1:** Primers used for Next-Generation Sequencing

| **Label** | | **Sequence** |
|---|---|---|
| Il23 | Il_for_ i5_506 | AATGATACGGCGACCACCGAGATCTACACACTGCATATCGTCGGCAGCGT CAGATGTGTATAAGAGACAGGCAGGAAGTGCTGAAGAAC |
| Il24 | Il_rev_ i7_706 | CAAGCAGAAGACGGCATACGAGATCATGCCTAGTCTCGTGGGCTCGGAGA TGTGTATAAGAGACAGGATGAAAAATTTCCTTAATGTTCTGTTCAATATG ATG |
| Il25 | Il_for_ i5_507 | AATGATACGGCGACCACCGAGATCTACACAAGGAGTATCGTCGGCAGCGT CAGATGTGTATAAGAGACAGGCAGGAAGTGCTGAAGAAC |
| Il26 | Il_rev_ i7_707 | CAAGCAGAAGACGGCATACGAGATGTAGAGAGGTCTCGTGGGCTCGGAGA TGTGTATAAGAGACAGGATGAAAAATTTCCTTAATGTTCTGTTCAATATG ATG |
| Il27 | Il_for_ i5_508 _ | AATGATACGGCGACCACCGAGATCTACACCTAAGCCTTCGTCGGCAGCGT CAGATGTGTATAAGAGACAGGCAGGAAGTGCTGAAGAAC |
| Il28 | Il_rev_ i7_708 | CAAGCAGAAGACGGCATACGAGATCCTCTCTGGTCTCGTGGGCTCGGAGA TGTGTATAAGAGACAGGATGAAAAATTTCCTTAATGTTCTGTTCAATATG ATG |

**Table S2**: Literature values for the uncatalyzed background hydrolysis rates of phosphodiester substrates

| $k_{uncat}$ ($s^{-1}$) | Substrate | Conditions | Reference |
|---|---|---|---|
| $2.6 \times 10^{-13}$ | *p*-nitrophenyl ethylphosphate | pH 7.5, 30 ˚C | [5] |
| $4.0 \times 10^{-14}$ | diphenyl phosphate | pH 7.5, 30 ˚C | [9,5] |
| $7 \times 10^{-16}$ | dineopentyl phosphate | pH 7, 25 ˚C | [7] |
| $3 \times 10^{-15\,a}$ | cAMP | pH 7, 25 ˚C | [6] |

4. $^{a}$ Extrapolated from $k_{uncat} = 3.0 \times 10^{-7}$ $s^{-1}$ for ethylene phosphate (a cyclic phosphodiester) at pH 7 and 100 ˚C.

**Table S3**: Overview of representative members from the three classes of cAMP-hydrolysing phosphodiesterases.

| Protein | Uniprot ID | Organism | Class | Superfamily | Metal requirement | $K_M$ (µM) | $k_{cat}$ ($s^{-1}$) | $k_{cat}/K_M$ ($M^{-1}$ $s^{-1}$) | Reference |
|---|---|---|---|---|---|---|---|---|---|

| | | | | | | | | | |
|---|---|---|---|---|---|---|---|---|---|
| PDE4C | Q08493 | *Homo sapiens* | I | HD-domain phosphodiesterase (Pfam CL0237) | Site 1: $Zn^{2+}$ Site 2: $Mg^{2+}$ or $Mn^{2+}$ | 1.7 | 0.41 | $2.4 \times 10^5$ | [10,11] |
| CpdP | Q56686 | *Aliivibrio fischeri* | II | Metallo-hydrolase/oxidoreductase (Pfam CL0381) | $Zn^{2+}$ (replacement with $Cu^{2+}$, $Mg^{2+}$, and $Ca^{2+}$ possible) | 73 | 2000 | $2.8 \times 10^7$ | [12,13] |
| CpdA | P0AEW4 | *Escherichia coli* | III | Calcineurin-like phosphoesterase (Pfam CL0163) | $Fe^{2+}$ | 500 | 1 | $2.0 \times 10^3$ | [4,14] |

**Table S4**: Primers for mutagenesis. Annealing regions are printed in black, mismatches are highlighted in red.

| Name | Sequence | $T_{annealing}$ (°C) |
|---|---|---|
| H20A_For | GCA AAA AAC GAC CGC AGC GGC | 55 |
| H20A_Rev | AAG GTT CTT CAG CAC TTC CTG CAG | |
| H31A_For | GCA GAT GTT GAT AAC CAT CTG CAG AAC GTG | 53 |
| H31A_Rev | AAT GTT ATC CTT GCC GCT GCG | |
| H36A_For | GCA CTG CAG AAC GTG ATT GAA GAT ATT CAT GAT TTT ATG | 53 |
| H36A_Rev | GTT ATC AAC ATC ATG AAT GTT ATC CTT GCC G | |
| H45A_For | GCA GAT TTT ATG CAG GCG GCG GC | 53 |
| H45A_Rev | AAT ATC TTC AAT CAC GTT CTG CAG ATG GTT ATC AAC | |
| C57A_For | GCG GCG GCA AAC GCA AGG AAA TGA | 55 |
| C57A_Rev | TGC CGC CGC CTG CAT AAA ATC ATG | |
| mini-cAMPase-Fix_For | GCG GCG GCA AAC TGC AGG AAA TG | 59 |

| | | |
|---|---|---|
| mini-cAMPase-Fix_Rev | TGC CGC CGC CCT GCA TAA AAT CAT G | |
| Short-824 _For | CGG TGG CAG CGG CGG CAAG | 63 |
| Short-824 _Rev | CTT GCA TGA AGT CGT GGA TGT CTT CGA TGA CGT TCT GCA AGT GG | |
| His Tag TEV mini-cAMPase For | ATG GGT TCT AGC CAC CAC CAC CAC CAC CAC TCT AGC GGT GAG AAT CTT TAT TTT CAG GGC ATG TAT GGC AAA CTG AAC GAT CTG | 50 |
| His Tag mini-cAMPase Rev | ATG GTA TAT CTC CTT CTT AAA GTT AAA CAA AAT TAT TTC | |

#### 5. SEQUENCES

Protein and nucleotide sequences are rendered in fasta format. For plasmid sequences, the coding region of the gene insert is highlighted in green.

>S-824 Protein
MYGKLNDLLEDLQEVLKNLHKNWHGGKDNLHDVDNHLQNVIEDIHDFMQGGGSGGKLQEMMK
EFQQVLDELNNHLQGGKHTVHHIEQNIKEIFHHLEELVHR*

>mini-cAMPase Protein
MYGKLNDLLEDLQEVLKNLHKNDRSGKDNIHDVDNHLQNVIEDIHDFMQAAAAAANCRK*

>pASK-IBA5plus-S824
GTGCTTTACTAAGTCATCGCGATGGAGCAAAAGTACATTTAGGTACACGGCCTACAGAAAAA
CAGTATGAAACTCTCGAAAATCAATTAGCCTTTTTATGCCAACAAGGTTTTTCACTAGAGAA
TGCATTATATGCACTCAGCGCAGTGGGGCATTTTACTTTAGGTTGCGTATTGGAAGATCAAG
AGCATCAAGTCGCTAAAGAAGAAAGGGAAACACCTACTACTGATAGTATGCCGCCATTATTA
CGACAAGCTATCGAATTATTTGATCACCAAGGTGCAGAGCCAGCCTTCTTATTCGGCCTTGA
ATTGATCATATGCGGATTAGAAAAACAACTTAAATGTGAAAGTGGGTCTTAAAAGCAGCATA
ACCTTTTTCCGTGATGGTAACTTCACTAGTTTAAAAGGATCTAGGTGAAGATCCTTTTTGAT
AATCTCATGACCAAAATCCCTTAACGTGAGTTTTCGTTCCACTGAGCGTCAGACCCCGTAGA
AAAGATCAAAGGATCTTCTTGAGATCCTTTTTTTCTGCGCGTAATCTGCTGCTTGCAAACAA
AAAAACCACCGCTACCAGCGGTGGTTTGTTTGCCGGATCAAGAGCTACCAACTCTTTTTCCG
AAGGTAACTGGCTTCAGCAGAGCGCAGATACCAAATACTGTCCTTCTAGTGTAGCCGTAGTT
AGGCCACCACTTCAAGAACTCTGTAGCACCGCCTACATACCTCGCTCTGCTAATCCTGTTAC
CAGTGGCTGCTGCCAGTGGCGATAAGTCGTGTCTTACCGGGTTGGACTCAAGACGATAGTTA
CCGGATAAGGCGCAGCGGTCGGGCTGAACGGGGGGTTCGTGCACACAGCCCAGCTTGGAGCG
AACGACCTACACCGAACTGAGATACCTACAGCGTGAGCTATGAGAAAGCGCCACGCTTCCCG
AAGGGAGAAAGGCGGACAGGTATCCGGTAAGCGGCAGGGTCGGAACAGGAGAGCGCACGAGG
GAGCTTCCAGGGGGAAACGCCTGGTATCTTTATAGTCCTGTCGGGTTTCGCCACCTCTGACT
TGAGCGTCGATTTTTGTGATGCTCGTCAGGGGGGCGGAGCCTATGGAAAAACGCCAGCAACG
CGGCCTTTTTACGGTTCCTGGCCTTTTGCTGGCCTTTTGCTCACATGACCCGACACCATCGA
ATGGCCAGATGATTAATTCCTAATTTTTGTTGACACTCTATCATTGATAGAGTTATTTTACC
ACTCCCTATCAGTGATAGAGAAAAGTGAAATGAATAGTTCGACAAAAATCTAGAAATAATTT
TGTTTAACTTTAAGAAGGAGATATACATATGTATGGCAAGTTGAACGACCTGCTGGAAGACT
TGCAAGAGGTGCTGAAGAACCTCCACAAAAACTGGCACGGTGGCAAAGACAACCTGCACGAC
GTCGACAACCACTTGCAGAACGTCATCGAAGACATCCACGACTTCATGCAAGGCGGTGGCAG
CGGCGGCAAGCTGCAAGAGATGATGAAAGAGTTCCAACAGGTGTTGGACGAACTCAACAACC
ACTTGCAAGGCGGTAAACACACCGTGCACCACATCGAACAAAACATCAAAGAGATCTTCCAC
CACTTGGAAGAGCTTGTACATCGCTAAGGATCCCTCGAGGTCGACCTGCAGGGGGACCATGG
TCTCTGATATCTAACTAAGCTTGACCTGTGAAGTGAAAAATGGCGCACATTGTGCGACATTT
TTTTTGTCTGCCGTTTACCGCTACTGCGTCACGGATCTCCACGCGCCCTGTAGCGGCGCATT
AAGCGCGGCGGGTGTGGTGGTTACGCGCAGCGTGACCGCTACACTTGCCAGCGCCCTAGCGC
CCGCTCCTTTCGCTTTCTTCCCTTCCTTTCTCGCCACGTTCGCCGGCTTTCCCCGTCAAGCT
CTAAATCGGGGGCTCCCTTTAGGGTTCCGATTTAGTGCTTTACGGCACCTCGACCCCAAAAA

ACTTGATTAGGGTGATGGTTCACGTAGTGGGCCATCGCCCTGATAGACGGTTTTTCGCCCTT
TGACGTTGGAGTCCACGTTCTTTAATAGTGGACTCTTGTTCCAAACTGGAACAACACTCAAC
CCTATCTCGGTCTATTCTTTTGATTTATAAGGGATTTTGCCGATTTCGGCCTATTGGTTAAA
AAATGAGCTGATTTAACAAAAATTTAACGCGAATTTTAACAAAATATTAACGCTTACAATTT
CAGGTGGCACTTTTCGGGGAAATGTGCGCGGAACCCCTATTTGTTTATTTTTCTAAATACAT
TCAAATATGTATCCGCTCATGAGACAATAACCCTGATAAATGCTTCAATAATATTGAAAAAG
GAAGAGTATGAGTATTCAACATTTCCGTGTCGCCCTTATTCCCTTTTTTGCGGCATTTTGCC
TTCCTGTTTTTGCTCACCCAGAAACGCTGGTGAAAGTAAAAGATGCTGAAGATCAGTTGGGT
GCACGAGTGGGTTACATCGAACTGGATCTCAACAGCGGTAAGATCCTTGAGAGTTTTCGCCC
CGAAGAACGTTTTCCAATGATGAGCACTTTTAAAGTTCTGCTATGTGGCGCGGTATTATCCC
GTATTGACGCCGGGCAAGAGCAACTCGGTCGCCGCATACACTATTCTCAGAATGACTTGGTT
GAGTACTCACCAGTCACAGAAAAGCATCTTACGGATGGCATGACAGTAAGAGAATTATGCAG
TGCTGCCATAACCATGAGTGATAACACTGCGGCCAACTTACTTCTGACAACGATCGGAGGAC
CGAAGGAGCTAACCGCTTTTTTGCACAACATGGGGGATCATGTAACTCGCCTTGATCGTTGG
GAACCGGAGCTGAATGAAGCCATACCAAACGACGAGCGTGACACCACGATGCCTGTAGCAAT
GGCAACAACGTTGCGCAAACTATTAACTGGCGAACTACTTACTCTAGCTTCCCGGCAACAAT
TGATAGACTGGATGGAGGCGGATAAAGTTGCAGGACCACTTCTGCGCTCGGCCCTTCCGGCT
GGCTGGTTTATTGCTGATAAATCTGGAGCCGGTGAGCGTGGCTCTCGCGGTATCATTGCAGC
ACTGGGGCCAGATGGTAAGCCCTCCCGTATCGTAGTTATCTACACGACGGGGAGTCAGGCAA
CTATGGATGAACGAAATAGACAGATCGCTGAGATAGGTGCCTCACTGATTAAGCATTGGTAG
GAATTAATGATGTCTCGTTTAGATAAAAGTAAAGTGATTAACAGCGCATTAGAGCTGCTTAA
TGAGGTCGGAATCGAAGGTTTAACAACCCGTAAACTCGCCCAGAAGCTAGGTGTAGAGCAGC
CTACATTGTATTGGCATGTAAAAAATAAGCGGGCTTTGCTCGACGCCTTAGCCATTGAGATG
TTAGATAGGCACCATACTCACTTTTGCCCTTTAGAAGGGGAAAGCTGGCAAGATTTTTTACG
TAATAACGCTAAAAGTTTTAGAT

>pASK-IBA5plus-library, with R = A/G, V = A/C/G, N = A/T/C/G, and D = A/G/T
TAATGAGGTCGGAATCGAAGGTTTAACAACCCGTAAACTCGCCCAGAAGCTAGGTGTAGAGC
AGCCTACATTGTATTGGCATGTAAAAAATAAGCGGGCTTTGCTCGACGCCTTAGCCATTGAG
ATGTTAGATAGGCACCATACTCACTTTTGCCCTTTAGAAGGGGAAAGCTGGCAAGATTTTTT
ACGTAATAACGCTAAAAGTTTTAGATGTGCTTTACTAAGTCATCGCGATGGAGCAAAAGTAC
ATTTAGGTACACGGCCTACAGAAAAACAGTATGAAACTCTCGAAAATCAATTAGCCTTTTTA
TGCCAACAAGGTTTTTCACTAGAGAATGCATTATATGCACTCAGCGCAGTGGGGCATTTTAC
TTTAGGTTGCGTATTGGAAGATCAAGAGCATCAAGTCGCTAAAGAAGAAAGGGAAACACCTA
CTACTGATAGTATGCCGCCATTATTACGACAAGCTATCGAATTATTTGATCACCAAGGTGCA
GAGCCAGCCTTCTTATTCGGCCTTGAATTGATCATATGCGGATTAGAAAAACAACTTAAATG
TGAAAGTGGGTCTTAAAAGCAGCATAACCTTTTTCCGTGATGGTAACTTCACTAGTTTAAAA
GGATCTAGGTGAAGATCCTTTTTGATAATCTCATGACCAAAATCCCTTAACGTGAGTTTTCG
TTCCACTGAGCGTCAGACCCCGTAGAAAAGATCAAAGGATCTTCTTGAGATCCTTTTTTTCT
GCGCGTAATCTGCTGCTTGCAAACAAAAAAACCACCGCTACCAGCGGTGGTTTGTTTGCCGG
ATCAAGAGCTACCAACTCTTTTTCCGAAGGTAACTGGCTTCAGCAGAGCGCAGATACCAAAT
ACTGTCCTTCTAGTGTAGCCGTAGTTAGGCCACCACTTCAAGAACTCTGTAGCACCGCCTAC
ATACCTCGCTCTGCTAATCCTGTTACCAGTGGCTGCTGCCAGTGGCGATAAGTCGTGTCTTA
CCGGGTTGGACTCAAGACGATAGTTACCGGATAAGGCGCAGCGGTCGGGCTGAACGGGGGGT
TCGTGCACACAGCCCAGCTTGGAGCGAACGACCTACACCGAACTGAGATACCTACAGCGTGA

GCTATGAGAAAGCGCCACGCTTCCCGAAGGGAGAAAGGCGGACAGGTATCCGGTAAGCGGCA
GGGTCGGAACAGGAGAGCGCACGAGGGAGCTTCCAGGGGGAAACGCCTGGTATCTTTATAGT
CCTGTCGGGTTTCGCCACCTCTGACTTGAGCGTCGATTTTTGTGATGCTCGTCAGGGGGGCG
GAGCCTATGGAAAAACGCCAGCAACGCGGCCTTTTTACGGTTCCTGGCCTTTTGCTGGCCTT
TTGCTCACATGACCCGACACCATCGAATGGCCAGATGATTAATTCCTAATTTTTGTTGACAC
TCTATCATTGATAGAGTTATTTTACCACTCCCTATCAGTGATAGAGAAAAGTGAAATGAATA
GTTCGACAAAAATCTAGAAATAATTTTGTTTAACTTTAAGAAGGAGATATACCATATGTATG
GCAAACTGAACGATCTGCTGGAAGATCTGCAGGAAGTGCTGAAGAAC**NDT**CATAAAAAC**VRC**
**VRCRRCRRC**AAGGATAAC**NDT**CATGAT**NDT**GATAACCATCTGCAGAACGTGATTGAAGATAT
TCATGATTTTATGCAGGGCGGCGGCAGCGGCGGCAAACTGCAGGAAATGATGAAAGAATTCC
AGCAGGTGCTGGATGAA**NDT**AACAAC**VRCVRCVRCRRCRR**CAAACAT**NDTNDT**CATCATATT
GAACAGAACATTAAGGAAATTTTTCATCATCTGGAAGAACTGGTGCATAGATAAGGATCCCT
CGAGGTCGACCTGCAGGGGGACCATGGTCTCTGATATCTAACTAAGCTTGACCTGTGAAGTG
AAAAATGGCGCACATTGTGCGACATTTTTTTTGTCTGCCGTTTACCGCTACTGCGTCACGGA
TCTCCACGCGCCCTGTAGCGGCGCATTAAGCGCGGCGGGTGTGGTGGTTACGCGCAGCGTGA
CCGCTACACTTGCCAGCGCCCTAGCGCCCGCTCCTTTCGCTTTCTTCCCTTCCTTTCTCGCC
ACGTTCGCCGGCTTTCCCCGTCAAGCTCTAAATCGGGGGCTCCCTTTAGGGTTCCGATTTAG
TGCTTTACGGCACCTCGACCCCAAAAAACTTGATTAGGGTGATGGTTCACGTAGTGGGCCAT
CGCCCTGATAGACGGTTTTTCGCCCTTTGACGTTGGAGTCCACGTTCTTTAATAGTGGACTC
TTGTTCCAAACTGGAACAACACTCAACCCTATCTCGGTCTATTCTTTTGATTTATAAGGGAT
TTTGCCGATTTCGGCCTATTGGTTAAAAAATGAGCTGATTTAACAAAAATTTAACGCGAATT
TTAACAAAATATTAACGCTTACAATTTCAGGTGGCACTTTTCGGGGAAATGTGCGCGGAACC
CCTATTTGTTTATTTTTCTAAATACATTCAAATATGTATCCGCTCATGAGACAATAACCCTG
ATAAATGCTTCAATAATATTGAAAAAGGAAGAGTATGAGTATTCAACATTTCCGTGTCGCCC
TTATTCCCTTTTTTGCGGCATTTTGCCTTCCTGTTTTTGCTCACCCAGAAACGCTGGTGAAA
GTAAAAGATGCTGAAGATCAGTTGGGTGCACGAGTGGGTTACATCGAACTGGATCTCAACAG
CGGTAAGATCCTTGAGAGTTTTCGCCCCGAAGAACGTTTTCCAATGATGAGCACTTTTAAAG
TTCTGCTATGTGGCGCGGTATTATCCCGTATTGACGCCGGGCAAGAGCAACTCGGTCGCCGC
ATACACTATTCTCAGAATGACTTGGTTGAGTACTCACCAGTCACAGAAAAGCATCTTACGGA
TGGCATGACAGTAAGAGAATTATGCAGTGCTGCCATAACCATGAGTGATAACACTGCGGCCA
ACTTACTTCTGACAACGATCGGAGGACCGAAGGAGCTAACCGCTTTTTTGCACAACATGGGG
GATCATGTAACTCGCCTTGATCGTTGGGAACCGGAGCTGAATGAAGCCATACCAAACGACGA
GCGTGACACCACGATGCCTGTAGCAATGGCAACAACGTTGCGCAAACTATTAACTGGCGAAC
TACTTACTCTAGCTTCCCGGCAACAATTGATAGACTGGATGGAGGCGGATAAAGTTGCAGGA
CCACTTCTGCGCTCGGCCCTTCCGGCTGGCTGGTTTATTGCTGATAAATCTGGAGCCGGTGA
GCGTGGCTCTCGCGGTATCATTGCAGCACTGGGGCCAGATGGTAAGCCCTCCCGTATCGTAG
TTATCTACACGACGGGGAGTCAGGCAACTATGGATGAACGAAATAGACAGATCGCTGAGATA
GGTGCCTCACTGATTAAGCATTGGTAGGAATTAATGATGTCTCGTTTAGATAAAAGTAAAGT
GATTAACAGCGCATTAGAGCTGCT

>pASK-IBA5plus-mini-cAMPase
TAATGAGGTCGGAATCGAAGGTTTAACAACCCGTAAACTCGCCCAGAAGCTAGGTGTAGAGC
AGCCTACATTGTATTGGCATGTAAAAAATAAGCGGGCTTTGCTCGACGCCTTAGCCATTGAG
ATGTTAGATAGGCACCATACTCACTTTTGCCCTTTAGAAGGGGAAAGCTGGCAAGATTTTTT
ACGTAATAACGCTAAAAGTTTTAGATGTGCTTTACTAAGTCATCGCGATGGAGCAAAAGTAC

ATTTAGGTACACGGCCTACAGAAAAACAGTATGAAACTCTCGAAAATCAATTAGCCTTTTTA
TGCCAACAAGGTTTTTCACTAGAGAATGCATTATATGCACTCAGCGCAGTGGGGCATTTTAC
TTTAGGTTGCGTATTGGAAGATCAAGAGCATCAAGTCGCTAAAGAAGAAAGGGAAACACCTA
CTACTGATAGTATGCCGCCATTATTACGACAAGCTATCGAATTATTTGATCACCAAGGTGCA
GAGCCAGCCTTCTTATTCGGCCTTGAATTGATCATATGCGGATTAGAAAAACAACTTAAATG
TGAAAGTGGGTCTTAAAAGCAGCATAACCTTTTTCCGTGATGGTAACTTCACTAGTTTAAAA
GGATCTAGGTGAAGATCCTTTTTGATAATCTCATGACCAAAATCCCTTAACGTGAGTTTTCG
TTCCACTGAGCGTCAGACCCCGTAGAAAAGATCAAAGGATCTTCTTGAGATCCTTTTTTTCT
GCGCGTAATCTGCTGCTTGCAAACAAAAAAACCACCGCTACCAGCGGTGGTTTGTTTGCCGG
ATCAAGAGCTACCAACTCTTTTTCCGAAGGTAACTGGCTTCAGCAGAGCGCAGATACCAAAT
ACTGTCCTTCTAGTGTAGCCGTAGTTAGGCCACCACTTCAAGAACTCTGTAGCACCGCCTAC
ATACCTCGCTCTGCTAATCCTGTTACCAGTGGCTGCTGCCAGTGGCGATAAGTCGTGTCTTA
CCGGGTTGGACTCAAGACGATAGTTACCGGATAAGGCGCAGCGGTCGGGCTGAACGGGGGGT
TCGTGCACACAGCCCAGCTTGGAGCGAACGACCTACACCGAACTGAGATACCTACAGCGTGA
GCTATGAGAAAGCGCCACGCTTCCCGAAGGGAGAAAGGCGGACAGGTATCCGGTAAGCGGCA
GGGTCGGAACAGGAGAGCGCACGAGGGAGCTTCCAGGGGGAAACGCCTGGTATCTTTATAGT
CCTGTCGGGTTTCGCCACCTCTGACTTGAGCGTCGATTTTTGTGATGCTCGTCAGGGGGGCG
GAGCCTATGGAAAAACGCCAGCAACGCGGCCTTTTTACGGTTCCTGGCCTTTTGCTGGCCTT
TTGCTCACATGACCCGACACCATCGAATGGCCAGATGATTAATTCCTAATTTTTGTTGACAC
TCTATCATTGATAGAGTTATTTTACCACTCCCTATCAGTGATAGAGAAAAGTGAAATGAATA
GTTCGACAAAAATCTAGAAATAATTTTGTTTAACTTTAAGAAGGAGATATACCATATGTATG
GCAAACTGAACGATCTGCTGGAAGATCTGCAGGAAGTGCTGAAGAACCTTCATAAAAACGAC
CGCAGCGGCAAGGATAACATTCATGATGTTGATAACCATCTGCAGAACGTGATTGAAGATAT
TCATGATTTTATGCAGGCGGCGGCAGCGGCGGCAAACTGCAGGAAATGATGAAAGAATTCCA
GCAGGTGCTGGATGAAGTTAACAACAGCGACCGCGGCGGCAAACATCATTTTCATCATATTG
AACAGAACATTAAGGAAATTTTTCATCATCTGGAAGAACTGGTGCATAGATAAGGATCCCTC
GAGGTCGACCTGCAGGGGGACCATGGTCTCTGATATCTAACTAAGCTTGACCTGTGAAGTGA
AAAATGGCGCACATTGTGCGACATTTTTTTTGTCTGCCGTTTACCGCTACTGCGTCACGGAT
CTCCACGCGCCCTGTAGCGGCGCATTAAGCGCGGCGGGTGTGGTGGTTACGCGCAGCGTGAC
CGCTACACTTGCCAGCGCCCTAGCGCCCGCTCCTTTCGCTTTCTTCCCTTCCTTTCTCGCCA
CGTTCGCCGGCTTTCCCCGTCAAGCTCTAAATCGGGGGCTCCCTTTAGGGTTCCGATTTAGT
GCTTTACGGCACCTCGACCCCAAAAAACTTGATTAGGGTGATGGTTCACGTAGTGGGCCATC
GCCCTGATAGACGGTTTTTCGCCCTTTGACGTTGGAGTCCACGTTCTTTAATAGTGGACTCT
TGTTCCAAACTGGAACAACACTCAACCCTATCTCGGTCTATTCTTTTGATTTATAAGGGATT
TTGCCGATTTCGGCCTATTGGTTAAAAAATGAGCTGATTTAACAAAAATTTAACGCGAATTT
TAACAAAATATTAACGCTTACAATTTCAGGTGGCACTTTTCGGGGAAATGTGCGCGGAACCC
CTATTTGTTTATTTTTCTAAATACATTCAAATATGTATCCGCTCATGAGACAATAACCCTGA
TAAATGCTTCAATAATATTGAAAAAGGAAGAGTATGAGTATTCAACATTTCCGTGTCGCCCT
TATTCCCTTTTTTGCGGCATTTTGCCTTCCTGTTTTTGCTCACCCAGAAACGCTGGTGAAAG
TAAAAGATGCTGAAGATCAGTTGGGTGCACGAGTGGGTTACATCGAACTGGATCTCAACAGC
GGTAAGATCCTTGAGAGTTTTCGCCCCGAAGAACGTTTTCCAATGATGAGCACTTTTAAAGT
TCTGCTATGTGGCGCGGTATTATCCCGTATTGACGCCGGGCAAGAGCAACTCGGTCGCCGCA
TACACTATTCTCAGAATGACTTGGTTGAGTACTCACCAGTCACAGAAAAGCATCTTACGGAT
GGCATGACAGTAAGAGAATTATGCAGTGCTGCCATAACCATGAGTGATAACACTGCGGCCAA
CTTACTTCTGACAACGATCGGAGGACCGAAGGAGCTAACCGCTTTTTTGCACAACATGGGGG

ATCATGTAACTCGCCTTGATCGTTGGGAACCGGAGCTGAATGAAGCCATACCAAACGACGAG
CGTGACACCACGATGCCTGTAGCAATGGCAACAACGTTGCGCAAACTATTAACTGGCGAACT
ACTTACTCTAGCTTCCCGGCAACAATTGATAGACTGGATGGAGGCGGATAAAGTTGCAGGAC
CACTTCTGCGCTCGGCCCTTCCGGCTGGCTGGTTTATTGCTGATAAATCTGGAGCCGGTGAG
CGTGGCTCTCGCGGTATCATTGCAGCACTGGGGCCAGATGGTAAGCCCTCCCGTATCGTAGT
TATCTACACGACGGGGAGTCAGGCAACTATGGATGAACGAAATAGACAGATCGCTGAGATAG
GTGCCTCACTGATTAAGCATTGGTAGGAATTAATGATGTCTCGTTTAGATAAAAGTAAAGTG
ATTAACAGCGCATTAGAGCTGCT

**Translation products of sequenced clones after Sorting 2 (Sanger sequencing)**

>1-A9
MYGKLNDLLEDLQEVLKNLHKNDRSGKDNIHDVDNHLQNVIEDIHDFMQAAAAAANCRK*

>1-B12
MYGKLNDLLEDLQEVLKNDHKNDSGGKDNIMILITICRT*

>1-D12
MYGKLNDLLEDLQEVLKNLHKNGSGGKDNLHDFDNHLQNVIEDIHDFMQGGGSGGKLQEMMK
EFQQVLDEVNNGGSGGKHFDHHIEQNIKEIFHHLEELVHR*

>1-F4
MYGKLNDLLEDLQEVLKNNHKNGGGGKDNFHDHDNHLQNVIEDIHDFMRAAAAAANCRK*

>2-A2
MYGKLNDLLEDLQEVLKNLHKNSGGSKDNVHDDDNHLQNVIEDIHDFMQAAAAAANCRK*

>2-A12
MYGKLNDLLEDLQEVLKNLHKNSGGNKDMILITICRT*

>2-B10
MYGKLNDLLEDLQEVLKNHHKNGDSGKDNVHDLDNHLQNVIEDIHDFMQGGGSGGKLRK*

>2-C8
MYGKLNYLLEDLQEVLKNIHKNDGGGKDNIMMVITICRT*

>2-E11
MYGKLNDLLEDLQEVLKIHKNDSGGKDNIHDDDNHLQNVIEDIHDFMQAAAAAANCRK*

>2-G3
MYGKLNDLLEDLQEVLKNHHKNGGGSKDNLHDHDNHLQNVIEDIHDFMQAAAAAANCRK*

>2-H1
MYGKLNDLLEDLQEVLKNFHKNGGSGKDNSHDLDNHLQNVIEDIHDFMQGGGSGGKLQEMMK
EFQQVLDELNNGGSSGKHISHHIEQNIKEIFHHLEELVHR*

>2-H4
MYGKLNDLLEDLQEVLKNLHKNGGDGKDNLHDRCRT*

>22-C2
MYGKLNDLLEDLQEVLKNYHKNDGGGKDNNHDIDNHLQNVIEDIHDFMQAAAAAANCRK*

>22-H8
MYGKLNDLLEDLQEVLKNIHKNDDSSKDNLHDRDNHLQNVIEDIHDFMQAAAAAANCRK*

#### SUPPLEMENTARY REFERENCES


1. Fischlechner, M. *et al.* Evolution of enzyme catalysts caged in biomimetic gel-shell beads. *Nat. Chem.* **6**, 791–796 (2014).
2. Check Hayden, E. Chemistry: Designer debacle. *Nature* **453**, 275–278 (2008).
3. O'Brien, P. J. & Herschlag, D. Functional Interrelationships in the Alkaline Phosphatase Superfamily: Phosphodiesterase Activity of *Escherichia coli* Alkaline Phosphatase. *Biochemistry* **40**, 5691–5699 (2001).
4. Imamura, R. *et al.* Identification of the cpdA Gene Encoding Cyclic 3′,5′-Adenosine Monophosphate Phosphodiesterase in Escherichia coli. *J. Biol. Chem.* **271**, 25423–25429 (1996).
5. van Loo, B. *et al.* An efficient, multiply promiscuous hydrolase in the alkaline phosphatase superfamily. *Proc. Natl. Acad. Sci.* **107**, 2740–2745 (2010).
6. Chin, J. & Zou, X. Catalytic hydrolysis of cAMP. *Can. J. Chem.* **65**, 1882–1884 (1987).
7. Schroeder, G. K., Lad, C., Wyman, P., Williams, N. H. & Wolfenden, R. The time required for water attack at the phosphorus atom of simple phosphodiesters and of DNA. *Proc. Natl. Acad. Sci.* **103**, 4052–4055 (2006).
8. Neun, S., Kaminski, T. S. & Hollfelder, F. Chapter Five - Single-cell activity screening in microfluidic droplets. in *Methods in Enzymology* (eds. Allbritton, N. L. & Kovarik, M. L.) vol. 628 95–112 (Academic Press, 2019).
9. Wolfenden, R., Ridgway, C. & Young, G. Spontaneous Hydrolysis of Ionized Phosphate Monoesters and Diesters and the Proficiencies of Phosphatases and Phosphodiesterases as Catalysts. *J. Am. Chem. Soc.* **120**, 833–834 (1998).
10. Wang, H. *et al.* Structures of the four subfamilies of phosphodiesterase-4 provide insight into the selectivity of their inhibitors. *Biochem. J.* **408**, 193–201 (2007).
11. Wang, P. *et al.* Expression, Purification, and Characterization of Human cAMP-Specific Phosphodiesterase (PDE4) Subtypes A, B, C, and D. *Biochem. Biophys. Res. Commun.* **234**, 320–324 (1997).
12. Dunlap, P. V. & Callahan, S. M. Characterization of a periplasmic 3':5'-cyclic nucleotide phosphodiesterase gene, cpdP, from the marine symbiotic bacterium Vibrio fischeri. *J. Bacteriol.* **175**, 4615–4624 (1993).
13. Callahan, S. M., Cornell, N. W. & Dunlap, P. V. Purification and Properties of Periplasmic 3':5'-Cyclic Nucleotide Phosphodiesterase. *J. Biol. Chem.* **270**, 17627–17632 (1995).
14. Richter, W. 3′,5′-Cyclic nucleotide phosphodiesterases class III: Members, structure, and catalytic mechanism. *Proteins Struct. Funct. Bioinforma.* **46**, 278–286 (2002).
